# Selection and Biodiversity change

**DOI:** 10.1101/527028

**Authors:** William Godsoe, Katherine E. Eisen, Daniel Stanton, Katherine M Sirianni

**Affiliations:** BioProtection Research Centre Lincoln University Lincoln New Zealand; Dept. of Ecology & Evolutionary Biology Cornell University Ithaca, NY. USA; Assistant Professor-Dept. of Ecology, Evolution and Behavior University of Minnesota Saint Paul, MN, USA; eCornell, Cornell University, Ithaca, NY, 14850, USA

**Keywords:** Price equation, theoretical ecology, evolutionary theory, biodiversity

## Abstract

There is a great need to understand how and why biodiversity, which we define as the variety of organisms found in a given place, changes over time. Current estimates suggest strikingly slow change in traditional measures of biodiversity. These estimates seem to contradict rapid shifts in the abundance of individual species and have led to a rethinking of the mechanisms shaping biodiversity. Conceptual models emphasize the role of competition among species or, more recently, selection on species identity (i.e. selection that favors some species at the expense of others). However, it is difficult to quantify how these mechanisms contribute to biodiversity change. To illustrate this point we present cases where strong competition or selection on species identity leads to no biodiversity change. In view of this disconnect we develop a new approach to studying biodiversity change using the Price equation. We show that biodiversity change responds to selection on species’ rarity, rather than to either competition or selection on species identity. We then show how this insight can be used to quantify the effects of the mechanisms previously thought to influence biodiversity: 1) selection, 2) (ecological) drift, 3) immigration and 4) speciation. Our results suggest the connection between species’ fates and their rarity is fundamental to understanding biodiversity change.

## Introduction

One of the fundamental goals of ecology is to understand how biodiversity changes over time (Tittensor et al. 2014). For our purposes, biodiversity is a summary of information about the variety of organisms found within a community (Vellend et al. 2017). Familiar indices of biodiversity include species richness, Gini-Simpson’s diversity, and Shannon entropy (Hill 1973; Jost 2007; Magurran 2013). Anthropogenic environmental changes including climate change, habitat destruction, and biological invasions pose major threats to biodiversity (Tittensor et al. 2014), but empirical measurements of change in traditional biodiversity metrics suggest a surprising degree of stasis, even when some species’ abundances shift dramatically (Dornelas et al. 2014; McGill et al. 2015; Vellend et al. 2013). This disconnect has led to a rethinking of the mechanisms in community ecology that can cause biodiversity to change (Chase and Knight 2013; Hillebrand et al. 2018).

Species interactions such as competition are thought to be a major driver of biodiversity change (MacArthur 1970; MacArthur 1972; Magurran 2013). For our purposes, species interactions occur when multiple species alter each other’s absolute finesses (i.e. their per capita growth rates) (Abrams 1987; Blanchet et al. 2020). Competition for example occurs when multiple species decrease each other’s per capita growth rates (Chase and Leibold 2003; Tilman 1980; Tilman and Wedin 1991). However, biodiversity—a summary of information about the variety of organisms found within a community—can be surprisingly insensitive to species interactions. Consider, for example, a community that consisted of 1 moose for every 2 squirrels 100 years ago (Figure 1 A). Competition for resources then leads to a drop in the squirrel population density so that the present community now consists of 2 moose for every 1 squirrel. Despite this competition biodiversity indices are unchanged: Simpsons remains 5/9, Shannon Diversity remains 0.637, species richness remains 2.

**Figure 1:**
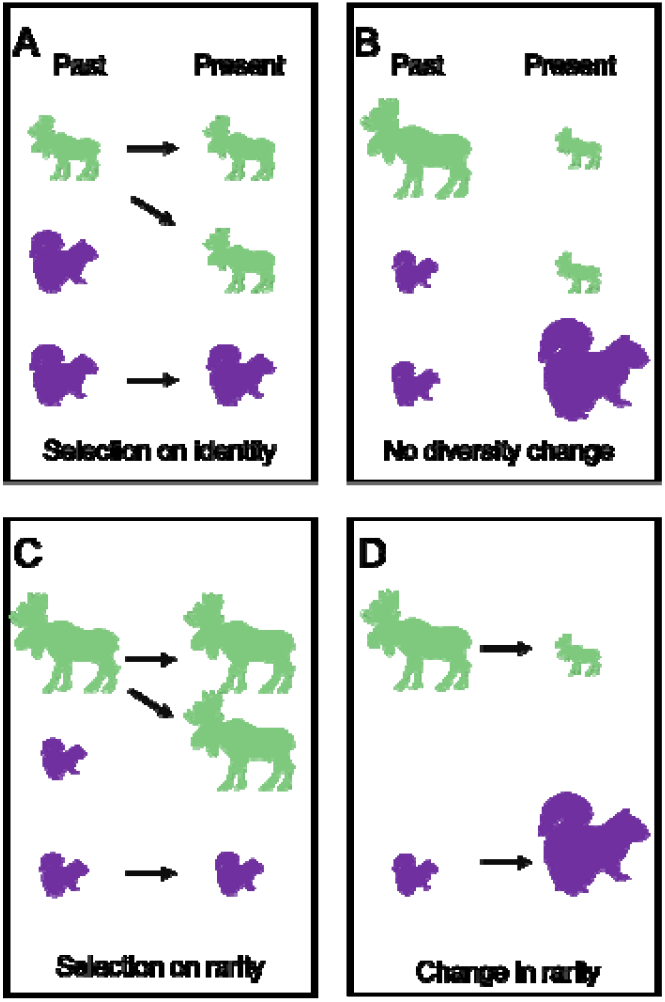
A) Selection on species identity results in each ancestral moose producing 2 descendants while each squirrel produces (on average) ½ a descendant and B) results in no diversity change. In this example, the size of each individual is scaled to its Shannon rarity scores: Moose past and 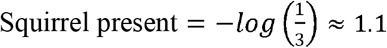; Moose present and Squirrel 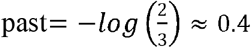. Shannon Diversity is the average of these rarity scores weighted by species’ frequency. In the past, diversity was: 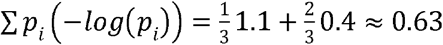. Diversity is unchanged in the present: 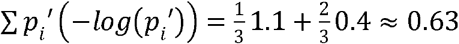. C) The lack of diversity change can be understood by focusing on the mechanisms that contribute to the changes in rarity scores. C) Selection acts in favour of the rare species (i.e., moose). All else being equal, this should increase the community’s average rarity and hence diversity. D) However, the effect of selection is opposed by changes in each species’ rarity scores. Moose were rare in the past, but their descendants are more common. Squirrels were common in the past, but their descendants are rarer. It is the balance between selection’s effects on rarity and change in rarity that produces net diversity change. Selection: 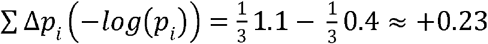 rarity changes: 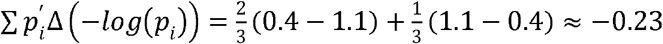.

To understand how biodiversity changes over time, there are alternatives to traditional analyses of species interactions. For example, diversity change might depend on species’ relative finesses (i.e. the frequency of descendants produced by each species). In view of this, many recent works discuss “selection” among species, i.e. change in species frequency due to differences in relative fitness. This use of the term “selection” emphasizes the distinction between absolute fitness and relative fitness. Analyses of selection are designed to study relative fitness (Rice 2004), while traditional measurements of species interactions produce a misleading picture of relative finesses (Abrams 1987), because these interactions inherently affect per-capita growth rates. Using the term selection in this way grounds analyses of biodiversity change in decades of work applying the notion of selection to the fates groups of organisms above the level of genotype, including species (Ayala and Campbell 1974; Barton and Servedio 2015; Day 2005; Jablonski 2008; Johnson and Stinchcombe 2007; MacArthur 1970; Norberg et al. 2012; Nowak 2006; Price 1970; Price 1995; Rabosky and McCune 2010; Rankin et al. 2015; Traulsen and Nowak 2006; Vellend 2010; Vellend 2016).

In particular, change in biodiversity may result from selection on species identity, the tendency of individuals belonging to some species to contribute more descendants than individuals belonging other species (Figure 1 A), averaged over many replicates to remove effects of chance (Jia et al. 2018; Perring et al. 2016; Purves and Turnbull 2010; Rosindell et al. 2011; Rosindell et al. 2012; Sherwin et al. 2017; Vellend 2010). The problems we illustrate with analyses of competition in Figure 1 apply to selection on species identity. This figure shows selection on species identity (i.e. individual moose contribute more descendants than do squirrels; Figure 1 A). But there is no biodiversity change.

We will argue that selection (among species) is the right lens to study biodiversity change, but only after we bridge a gap between two concepts: species identity and species’ rarity. Diversity indices are designed to be indifferent to information on species identity. Thus, when studying diversity, it matters that 1/3 individuals belong to one species, but it does not matter whether that species is a moose or a squirrel (Figure 1 B). Instead, diversity indices are designed to measure the “rarity” of species, averaged across a community (Hill 1973; Jost 2007; Patil and Taillie 1982). There is some latitude on how the term rarity should be interpreted. Rarity is always a function of species relative abundance (i.e. frequency), such that infrequent species are rarer than frequent ones. However, some measures of rarity heavily emphasize infrequent species (the definition used when studying species richness). Other measures of rarity place more emphasis on frequent species, such as Simpson’s diversity (Jost 2006; Jost 2007). This information on how diversity is measured is obscured when we study selection on species identity. One way to better understand diversity change is to consider selection’s effects on rarity explicitly.

Information on rarity can be incorporated in a more general definition of selection: the association between fitness of each type of organism and the measurement studied (Lehtonen 2018; Price 1970; Price 1972; Queller 2017; Rice 2004). Here fitness is a measure of frequency of descendants produced by each type of organism (which in turn depends on births and deaths for that species). Measurements represent an attribute of each type of organism which can be quantified (Frank 2012b; Price 1995). This interpretation of selection is used in many branches of evolutionary theory, notably quantitative genetics (Lande 1979; Lande and Arnold 1983) and its generalization, the Price equation (Price 1970; Queller 2017; Rice 2004). Ecologists have used this interpretation to study several attributes of ecological communities including body size, crop yield, and disease resistance (Collins and Gardner 2009; Genung et al. 2011; Govaert et al. 2016; Norberg et al. 2012). This interpretation of selection has also been used to study how the productivity of a community changes when biodiversity has been manipulated (Loreau and Hector 2001), but ecologists have not yet used this approach to study how biodiversity itself changes.

When the Price equation is used to partition biodiversity change under simplified conditions, two mechanisms emerge as fundamental. One mechanism is selection on rarity, which describes the fitness advantages of individuals belonging to rare species, although this does not preclude the possibility for selection on other traits. The other mechanism is rarity changes (more precisely, a weighted average of the changes in species’ rarities between past and present). Rarity changes represent a special case of the mechanism in evolutionary theory known as “transmission bias”, which describes how the measurements associated with each type of organism change over time (Frank 2012a; Frank 2012b). Using this insight, we show that the interplay between these two mechanisms explains why strong selection sometimes leads to zero biodiversity change. We then use simulations and empirical datasets to show how, under more ecologically relevant conditions, the effects of (ecological) drift, immigration, and speciation can be quantified with extensions of the Price equation. Together, these results show that change in biodiversity is linked to species rarity and that tools from evolutionary theory provide a natural framework to capture this dependency.

## The model

### Selection and biodiversity change

A natural way to develop intuition about evolution is to track the individual organisms’ contributions to change in the variable of interest (Frank 2012b; Kerr and Godfrev□Smith 2009). But what is an individual’s contribution to diversity? The usual way to express diversity indices focuses on the contributions of species, not the contributions of individuals. For example, consider the Shannon Weiner diversity index:

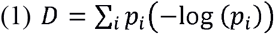

Where *p_i_* is the frequency of species i (the number of individuals belonging to species divided by the total number of individuals in the community). This quantity is also called relative abundance or proportional abundance. In turn, species’ *i* contribution to diversity (*p_i_*(−log (*p_i_*))) cleaves into two distinct parts. One component is the weighting factor *p_i_* which leads to species with more individuals contributing more to *D*. The other component is an individual’s contribution (−log (*p_i_*)) to *D*. This component is one way to measure the rarity of each individual in the community. In evolutionary biology, an individual’s contribution to D would be called a trait (Conner and Hartl 2004; Frank 2012b; Lande 1979; Lande and Arnold 1983). The weighting factor *p_i_* and the individual’s contributions (−log (*p_i_*)) to *D* serve different roles in Shannon Weiner entropy, but this fact is obscured because one measurement is a function of the other.

The distinction between individuals’ contributions and a weighting factor is used to interpret Shannon Weiner in other disciplines (Cover and Thomas 2012; Frank 2012c). We present this interpretation here, but readers already convinced by the distinction may skip this paragraph. Shannon Weiner entropy is interpreted as the average number of yes/no questions (or bits) needed to efficiently identify the category to which an individual observation belongs. In terms more familiar to an ecologist, this is the average number of couplets in an efficient dichotomous key needed to identify an individual to species (Jost 2006; Jost 2007). The number of questions needed to identify an individual with frequency *p_i_* is calculated by computing (−log_2_(*p_i_*)). Note the choice to compute logarithms in base 2, which highlights the connection to bits. For example, when an individual belongs to a community of two species where ½ of the individuals belong to species A, we can ask a single question to distinguish A from B corresponding to −log_2_ (*p_i_*)=1. When an individual belongs to a community where ¼ of individuals belong to each of four species A, B, C, D, then we can use two questions to efficiently identify an individual. For an individual belonging to A, the first question could be “is this individual a member of species A or B” with the next question being: “is this individual a member of species A”. These two questions correspond to −log_2_ (*p_i_*)=2. The interpretation of Shannon Weiner as a measure of the number of questions needed to identify individuals is possible because we distinguish unique contributions that −log (*p_i_*) and *p_i_* play. Some introductions to the concept of Shannon Wiener entropy go so far as to physically represent the number of questions we ask about an individual, see for example (Kahn Academy Labs 2014), which represents the number of questions asked about an individual as the number “bounces” a ball makes on pegs arranged to form a pachinko machine. Ironically, the pachinko machine will be familiar to some readers from long running CBS program “The Price is Right”. This video then uses equations to compute the frequency of different types of balls to compute overall entropy, reinforcing the distinct roles that −log (*p_i_*) and *p_i_* play in calculations of Shannon Wiener entropy.

Other diversity indices cleave into an individual’s rarity and a weighting factor (Hill 1973; Patil and Taillie 1982):

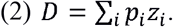

Examples are listed in (Table 1) including species richness, Simpson’s diversity and Gini-Simpson diversity. Additional diversity measures are functions of Equation 2, notably the diversity of an equivalent number of uniformly distributed species i.e., Hill numbers (Hill 1973, Jost 2007).

**Table 1.** Examples of diversity measures that reflect averages of rarity measures. Several such measures in include the exponent *q* which describes how sensitive a rarity measure is to changes in species’ frequency.

| Index | Rarity measure ($z_i$) | Expression |
| --- | --- | --- |
| Species Richness | $\frac{1}{p_i}$ | $\sum p_i z_i$ |
| Shannon Entropy | $-\log(p_i)$ | $\sum p_i z_i$ |
| Simpson's concentration | $p_i$ | $\sum p_i z_i$ |
| Gini-Simpson's index | $p_i$ | $1 - \sum p_i z_i$ |
| HCDT entropy | $p_i^{q-1}$ | $\frac{1}{q-1} \left(1 - \sum p_i z_i\right)$ |
| Reyni entropy | $p_i^{q-1}$ | $\frac{1}{q-1} \left(-\log \sum p_i z_i\right)$ |
| Hill numbers | $p_i^{q-1}$ | $\left(\sum p_i z_i\right)^{1/(1-q)}$ |

The distinction between frequency and rarity has been made in the ecological literature, but it is less prominent. For example, in a synthesis of disparate diversity indices, Hill (1973) emphasizes the distinct role that frequency plays by labelling frequency *p_i_* when it is used to compute rarity, then relabelling frequency *w_i_* when it is used as a weighting factor. Similarly, Patil and Taillie (1982) emphasize the distinction between species *i*’s frequency (which they denote *π_i_*) and its rarity (which they denote *R*(*i,π_i_*)). These authors take for granted that frequency and rarity are distinct concepts making unique contributions to diversity indices.

Diversity change is the difference between *D* in one time step and *D* in a subsequent time step:

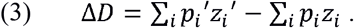

The past frequency of species *i* is *p_i_* and the present frequency is *p_i_*′. Similarly, past measurement of rarity is *z_i_* and present measurement of rarity is *z_i_*′. The difference is denoted Δ. Diversity reaches a maximum when all species are equally rare, then tends to decrease as species frequencies become less even. This definition of diversity change works over any timescale. By the same token, the definitions of selection and other mechanisms work over any timescale, although the importance of each mechanism can vary across timescales (Frank 2012a).

We propose that change in *D* can should be partitioned using the Price equation from evolutionary theory (Price 1970). To our knowledge the equation has not been applied to diversity indices, but it is used to partition many other averages that take the form of Equation 2 (Frank 2012b). In a community that is “closed” so that all individuals in the present community are descended from individuals of the past community. Equation 2 can be re-arranged into:

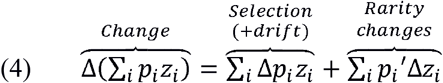

For a proof, see derivation of Equation 1 in Frank (2012b). Note that this division into two terms recapitulates the distinction between frequency change and rarity change. The first term measures the consequences of changes in species’ frequencies 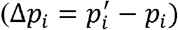. The second term describes consequences of changes in rarity 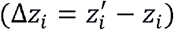 averaged across species.

In models, the first term of Equation 4 is typically interpreted as the result of selection (Price 1970). In nature, however, species frequencies change because of two interlinked causes: differences in the fitness of each species (i.e., selection) and sampling variation in the number of descendants produced by each species (i.e., drift). Both processes increase D when individuals belonging to rare species produce more offspring than individuals belonging to common species, and both processes decrease D when individuals belonging. As a result, it is easy to measure the joint effects of drift and selection, but it is difficult to tease apart their individual contributions Rice (2004). Over numerous replicated experiments, drift could be identified, in principle, as cases in which all species have the same fitness, averaged across many replicates and regardless of their frequency (see Engen and Sæther (2014) for a mathematical distinction).

Selection and drift act on species’ rarity in the past, and there is no guarantee that rarity will remain constant over time. This complication is captured by the second term in Equation 4, rarity changes. This term is large when a species’ rarity in the past differs from its rarity score in the present (Figure 1 D). This change in rarity explains why strong selection can lead to zero change in biodiversity. In the example in our introduction, selection favoured moose so strongly that moose and squirrel exchange rarity scores (Figure 1 D). Because these effects cancel out precisely, there is no net change. Species richness (one of the most common measures of diversity) is particularly sensitive to this problem. The species richness index defines rarity as 1/*p_i_* (Hill 1973; Jost 2006; Jost 2007). Substituting this definition into Equation 2, rarity changes precisely counterbalances selection, so long as all species that occurred in the past leave descendants (Appendix S1.1). However, when strong selection causes species to exchange frequencies as in our example (Figure 1 A), there will be no net change in diversity regardless of the index used (Appendix S1.2).

It is worth pausing here for a moment to think about why rarity changes must be considered when describing change in biodiversity. Selection changes species frequencies between the past and the present. Selection in the past acted on species’ rarity in the past. Selection in the past did not act on species’ rarity in the present (selection is powerful, but not psychic). In contrast, biodiversity change depends on the present rarity of each species. Thus, there is a gap between the attributes upon which selection can act (i.e., species’ rarity in the past) and biodiversity change (which depends on species’ rarity in the present). What Equation 4 shows is that rarity changes fill this gap.

### Partitioning biodiversity changes into multiple mechanisms

The derivation in the previous section assumes that all individuals in the present community descended from past community members. To allow for the arrival of new individuals we use the extended Price equation (Frank 2012b; Kerr and Godfrey□Smith 2009; Rice 2004). Selection and drift measure the number of descendants produced by each species. As a result, it is necessary to distinguish members of the present community that descend from the past community (with frequency *ω*), and new arrivals. To facilitate comparisons with Vellend (2016), we further divide new arrivals into two categories immigrants (with frequency *μ*), and individuals belonging to new species (with frequency *σ*). Following the arguments laid out in Vellend (2016), we assume that all individuals belong to one of these three categories such that *ω + μ + σ* = 1. Diversity in the present community can be expressed as:

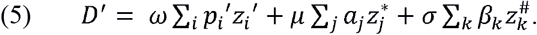

In Equation (5) diversity is divided into the contributions of individuals that are descendants’ immigrants and members of new species. Each individual will belong to only one category, but a single species may include some individuals that are immigrants and some individuals that are descendants of the past community. *p_i_*′ now represents the frequency of species *i* among descendants, and *a_j_* is the frequency of immigrants that belong to species *j* among all immigrants. *β_k_* is the frequency of individuals that belong to new species *k* among all individuals belonging to new species. The rarity of the *i^th^* descendant species is *z_i_*′, the rarity of the *j*^th^ immigrant species is 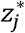, and the rarity of the k^th^ new species is 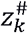. When a species includes descendants and immigrants 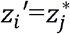.

Change in diversity can be partitioned with the extended Price equation (Kerr and Godfrey-Smith 2009). We use a derivation of this extension of the Price equation presented on pages 1014-105 of Frank (2012b), but we make one minor modification: new arrivals are divided into two categories, immigrants and new species. This leaves us with:

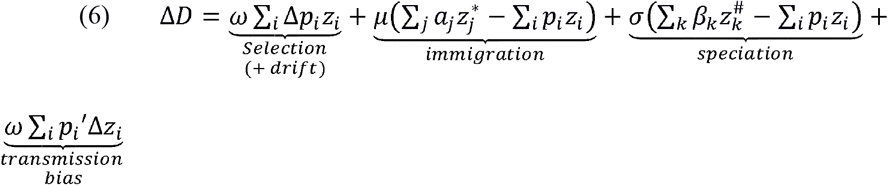

The selection + drift term is analogous to the selection term in Equation 4—it still describes the difference between the frequency of individuals in species *i* in the past and the frequency of the descendants associated with species *i* in the present (Δ*p_i_*). The effects of immigration depend on the proportion of individuals in the present community that are immigrants (*μ*), and the difference between the average rarity of immigrants in the present and the diversity of the past community. Similarly, the effects of speciation depend on the proportion of individuals that belong to new species (*σ*), the difference between the average rarity of new species in the present, and the average rarity of the past community. Essentially, we treat species identity in the way described by Vellend (2010): “Species identity is a categorical phenotype assumed to have perfect heritability, except when speciation occurs, after which new species identities are assigned.” With the extended Price equation, a natural way to assign new identities is to treat members of new species as new arrivals to the community.

The arrival of new species via immigration and speciation can indirectly increase the rarity of descendant species. This form of rarity change concerns the proportion of the community whose ancestors were present in the community in the past time step (*ω*). Change in species *i*′s rarity depends on the proportion of descendants belonging to species *i* (*p_i_*′), and the change in the rarity of species *i* (Δ*z_i_*).

### Empirical partitioning of biodiversity change

Our approach can be applied to partition empirical measurements of change in diversity. For example, (Remus-Emsermann et al. 2018) measured the frequency of two bacterial species (*Escherichia coli* and *Pantoea eucalypti*) on leaves of mouse ear cress (*Arabidopsis thaliana*). Leaves were inoculated to produce a population density of 2.5×10^6^ colony forming units of *E. coli* and 4.95×10^6^ colony forming units of *P. eucalypti* per 1 gram of tissue of *A. thaliana* (fresh weight). The population density was subsequently measured by assaying the number of colony forming units at 1, 3, and 7 days post-infection. The experiment was established to assay the frequency with which *E. coli* obtain plasmids from *P. eucalypti* via conjugation, but conjugation events were not observed in this treatment and so are not considered here (Remus-Emsermann et al. 2018). No speciation was observed, and the experimental protocol was designed to exclude immigration. Due to the large population sizes in this experiment, we expect selection to contribute to biodiversity change far more than drift. Figure 2 partitions the resulting diversity change. Initially diversity changes dramatically, indicating both selection against the rarer species (*E. coli*) and changes in rarity scores notably – *P. eucalypti* is far more common at the end of day 1 than it was at day 0. Later sampling intervals show modest selection and far weaker changes in rarity.

**Figure 2.**
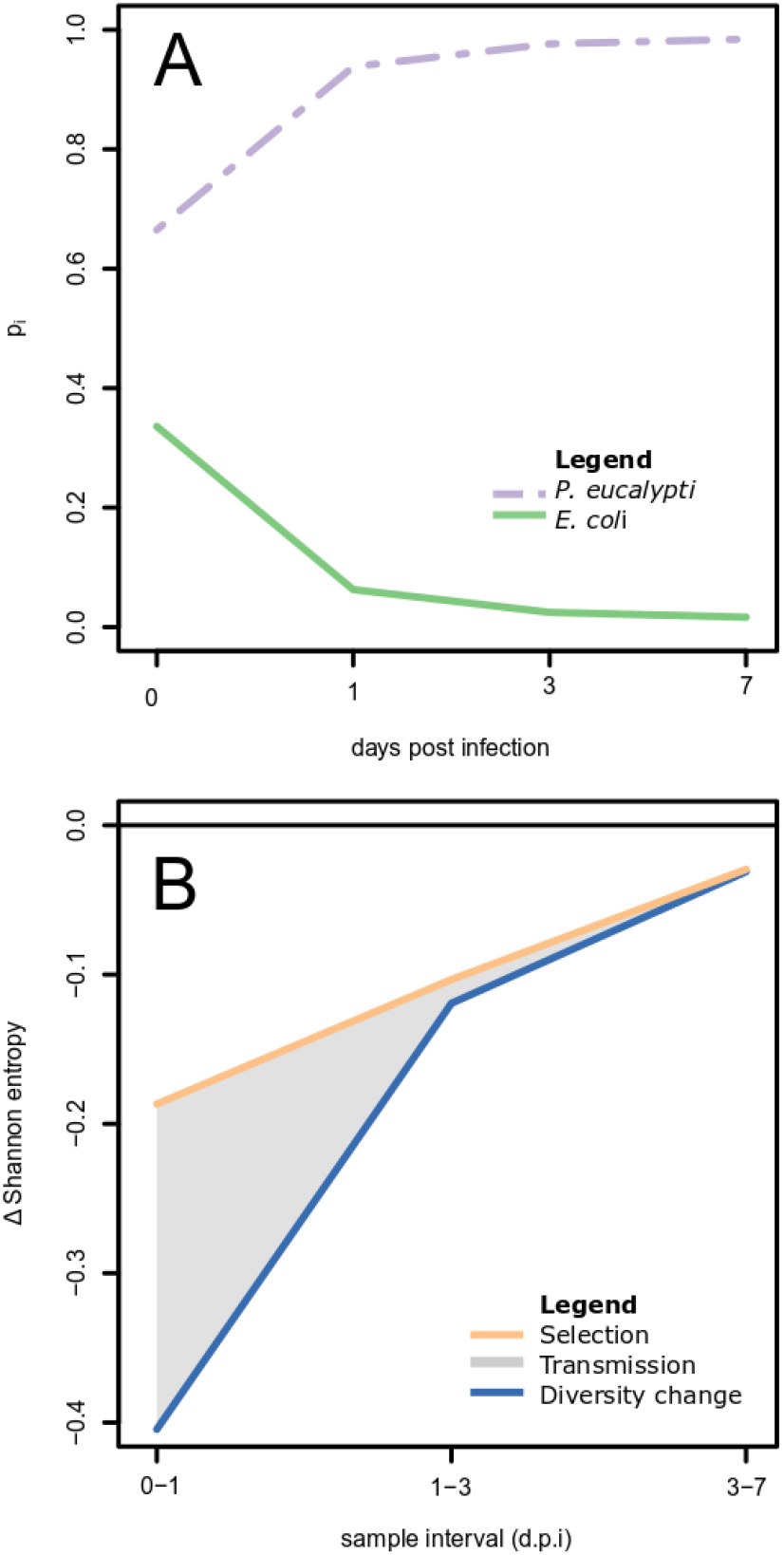
A) Mechanisms changing biodiversity in an experiment concerning two species of bacteria (*P. ecalypti*, purple; *E. coli*, green). Leaves of *A. thaliana* were inoculated with both species and the frequency of each species was measured at days 0, 1, 3, and 7, resulting in three sampling intervals during which we can measure change in biodiversity (0-1, 1-3 and 3-7 days post-inoculation). Over this time, the proportion of *E. coli* decreased, suggesting strong selection against this species. Since the species that was initially rarer (*E. coli*) decreased in frequency, Shannon-Wiener diversity decreases as well. Using Equation 4 total change in diversity partitions into the effect of selection on rarity (yellow) plus changes in rarity (grey shaded region). Change is rapid in the first sampling period resulting in contributions from both mechanisms. During later sampling periods change in rarity is less important.

When used in multispecies communities, our approach can help to identify and interpret gaps between selection on species identity and diversity change. For example, Sirianni (2018) tracked the abundance of water fleas (Cladocerans) that inhabit a rock pool on Appledore Island in the Gulf of Maine in the United States for 13 weeks over the summer of 2013. Generation times of many species within this community are short (Sirianni 2018), resulting in substantial fluctuations in biodiversity over the course of the summer. For example, individuals of the common species *B. rubens* can reproduce in as little as 1-2 days after birth in a lab environment (Morales-Ventura et al. 2018). Each week, Sirianni (2018) sampled 250 mL of a rock pool that averaged 250 L, and counted the zooplankton present in the sample to estimate density/L of each zooplankton species. Due to the large size of the communities, we expect selection to contribute to biodiversity change far more than drift, particularly for the abundant species (described below).

Although species frequencies fluctuated rapidly over the course of the summer, only some fluctuations substantially changed diversity. For example, there was a modest increase in diversity between weeks four and five (Figure 3 grey band), and a sharper decrease in diversity between weeks ten and eleven. This change is superficially puzzling since both periods show strong selection on species identity (Figure 3 B), notably selection in favor of *Brachionus rubens* (Figure 3 B Green).

**Figure 3.**
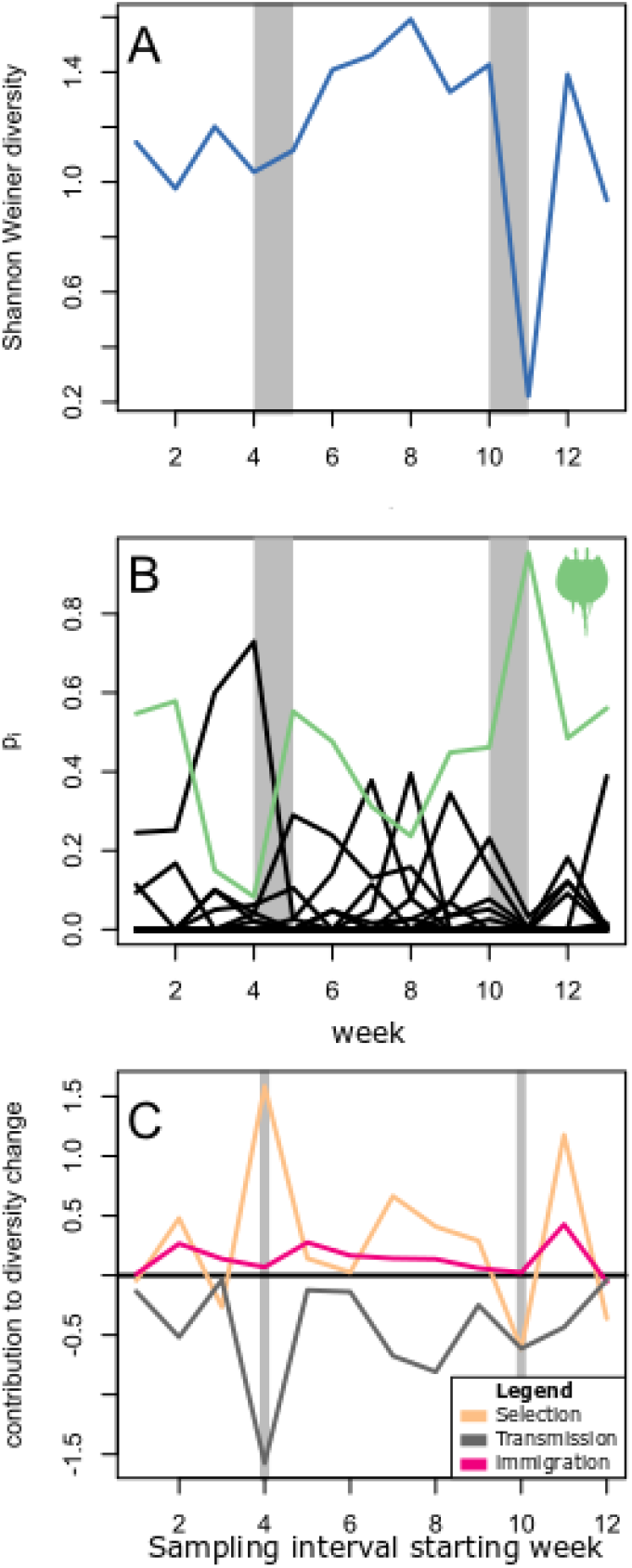
Shannon Wiener diversity change in species inhabiting a rock pool in Appledore Island. A) Over the course of 13 weeks, diversity changed gradually, with a sudden dip between weeks 10 and 11. The grey bars denote two sampling intervals discussed in the main text. B) Fluctuations of the frequencies of all species (black) with one abundant species highlighted (*B. rubens*, green). C) Partitioning of biodiversity change into contributions of selection or drift (dark grey), rarity changes (tan), and the arrival of new taxa (red), which in this case probably indicates taxa emerging from eggs.

The interplay of selection on rarity and changes in rarity better accounts for the change in diversity (Figure 3 C). Between weeks four and five, selection disfavored rare species, which strongly decreased diversity (dark grey). However, this effect was opposed by changes in species’ rarity (Figure 3 C yellow). Rare species became common (*Brachionus rubens*), and common species became rare (*Moina macrocopa*). In contrast, between weeks ten and eleven, all rare species became rarer while the most common species (*Brachionus rubens*) became vastly more common. As a result, both selection and rarity changes decreased diversity, leading to a large net decrease in diversity between weeks ten and eleven as opposed to the modest increase between weeks four and five. The arrival of new species increased diversity slightly throughout the sampling period (Figure 3 C red).

### Simulated changes including multiple mechanisms

Though the empirical examples above show how selection on rarity and other mechanisms can be measured, they offer limited opportunities to contrast our approach with others. To compare our models with existing analyses of selection and species interactions, we use simulations [see Online Appendix S2 for R scripts (R_Core_Team)]. All scripts were initially based on examples provided by (MacDonald and Vellend 2016). Though most of these examples have been heavily modified. The majority of the simulations consider competition based on the Ricker model (Luís et al. 2011; Otto and Day 2007; Ricker 1954). A model related to Lotka-Volterra competition but in discrete time and hence easier to compare to Equations 4 and 6:

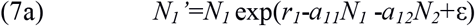

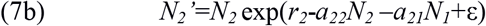

*N_i_* denotes the number of individuals of species *i* in the previous generation and *N_i_’* denotes the number of individuals in the current generation. *N_i_’* depends on the number of individuals of that species at time *t* multiplied by a term describing population growth. The *r_i_* parameters describe the intrinsic growth rates of species *i*. The *a_ij_* terms describe how interacting with species *j* decreases the growth rate of species *i*. These terms represent interspecific interactions when *i* and *j* differ and intraspecific interactions when *i* and *j* are the same. We have included stochasticity in only our model of drift by adding a noise term ε that is normally distributed with a mean of 0 and a standard deviation of 0.02. This is a model of absolute fitness (i.e. change species’ abundances). Data on absolute fitness can in turn be used to compute change in species’ frequencies (Mallet 2012), which is used, in turn, to derive the terms in Equation 4. In addition, we compute the effects of selection or drift on species identity (Appendix S1.3 for details).

Equation 7 results in cases where some species thrive in the face of competition. This results in individuals belonging to one species producing more descendants than individuals belonging to other species (i.e. selection among species). This is easiest to illustrate in discrete models where ε=0. One scenario is frequency-independent selection in favor of species 1. This occurs when individuals belonging to species 1 invariably produce more descendants. To simulate this, we this set species 1’s intrinsic growth rate to be higher than species 2’s and set all competition coefficients are equal (scenario 1: *a_11_= a_22_=a_12_=a_21_*=0.001, *r_1_*=0.6, *r_2_*=0.3). Another possibility is frequency-dependent selection where individuals of a given species have an advantage when that species is rare. To simulate this we set interspecific competition coefficients to be less than intraspecific competition coefficients (Scenario 2: *a_12_=a_21_*=0.01, *a_11_= a_22_*=0.001*r_1_*=0.6, *r_2_*=0.3; Mallet 2012).

Drift occurs when neither species has an advantage and there are stochastic fluctuations in species frequency (i.e. where ε>0). To simulate drift (scenario 3), we set the intrinsic growth rates of the two species to be equal (*r_1_*=0.3, *r_2_*=0.3) and set all competition coefficients to be equal (*a_11_= a_22_=a_12_=a_21_*=0.001; Adler et al. 2007).

To consider the consequences of immigration into a patch (scenario 4), we modelled dispersal among a metacommunity of three patches (labelled *L*=1,2,3). To do this, we divided the life cycle of each species into two stages. The first stage represents local population growth in a patch using equations Equation 7a and 7b (with an added index to indicate patch number). In this simulation interspecific competition was weaker than intraspecific competition (*a_12_=a_21_*=0.01, *a_11_= a_22_*=0.001). Species 1 had a lower intrinsic growth rate than species 2 in patch 1 and a higher growth rate than species 2 in patch 3. In patch 2, intrinsic growth rates were the same for both species (*r_11_*=0.3, *r_21_*=0.9, *r_12_*=0.6, *r_22_*=0.6, *r_13_*=0.9, *r_23_*=0.3). The density of species *i* after local population growth is denoted *N_iL_^*^*. The second stage describe dispersal among patches (Hedrick 2011):

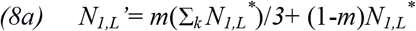

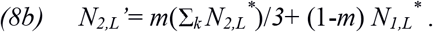

In this community, a proportion *m=0.05* of individuals joined a regional species pool. An equal proportion (1/3) of these individuals moved to each patch from the regional species pool. A proportion (1-*m*) of individuals remain in their birth patch (i.e., they do not immigrate). [see Online Appendix S2 for R scripts (R_Core_Team)]. All scripts were initially based on examples provided by (MacDonald and Vellend 2016). Though most of these examples have been heavily modified.

Lastly, we examined speciation (scenario 5) with a variant of Hubbell (2001)’s neutral model where biodiversity is governed by drift and speciation. This model is not based on the Ricker equation but instead consists of a fixed metacommunity of *j* individuals. At each time step, individuals die with some probability. They are replaced by the descendants of another individual selected at random from other individuals in the community (representing drift) or by a new species (representing speciation).

In closed communities, biodiversity change is tightly linked to selection on rarity and change in rarity. In a simulation with frequency independent selection (scenario 1), the favored species (species 1 in green, Figure 4 A) is initially rarer than the other species (species 2 in purple, Figure 4 A). However, by the end of the simulation, species 1 is far more common. As a result, biodiversity initially increases and then declines (Figure 4 B; blue). Selection on species identity is always positive, such that it fails to reproduce this pattern in diversity change (Figure 4 B; yellow). In contrast, selection on species rarity (Figure 4 C; yellow) closely matches biodiversity change, with a small discrepancy because of rarity changes (Figure 4 C; grey). A simulation of negative frequency-dependent selection (scenario 2) leads to coexistence between two species (Figure 4 D). Change in biodiversity is positive throughout the simulation (Figure 4 E; blue), but peaks sooner than selection on species identity (Figure 4 E; yellow). In contrast, selection on rarity (Figure 4 F; yellow) closely tracks biodiversity change (Figure 4 F; blue) with a small discrepancy because of rarity changes (Figure 4 F; grey). Under a simulation of drift (scenario 3; Figure 4 G), the frequency of species 1 changes in a manner analogous to selection on species identity (Figure 4 H), but selection on species identity for species 1 (Figure 4 H; yellow) is far larger than biodiversity change (Figure 4 H; blue). In this simulation, biodiversity change is equal to the combined effects of drift and rarity changes (Figure 4 I).

**Figure 4.**
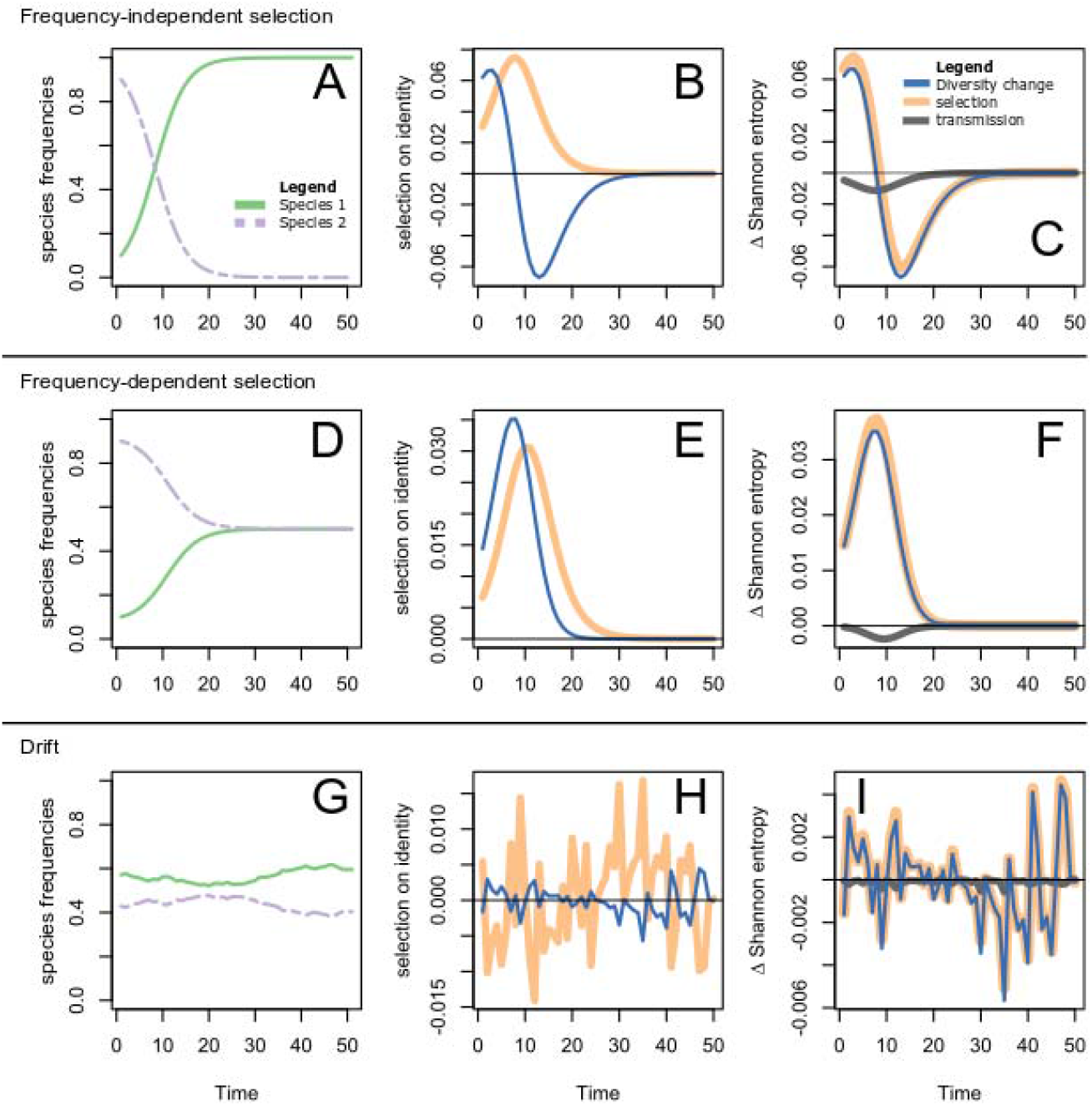
Comparison of selection and biodiversity change. A) Under frequency-independent selection, the frequency of species 1 (green) increases at the expense of species 2 (purple). B) Selection on species identity (See Online appendix S1.3) is always positive, indicating that species 1 is always fitter than species 2 (yellow). In contrast Shannon Wiener diversity increases when species 1 is rare then decreases when species 1 is common (blue). C) Selection on rarity (yellow; measured using Equation 4) shows the same trend as biodiversity change (blue), increasing when species 1 is rare and decreasing when species 1 is common. Rarity changes (grey; measured using Equation 4) accounts for the discrepancy between selection and biodiversity change. D) Frequencies of two species under negative frequency-dependent selection leading to coexistence (scenario 2). E) In this case, biodiversity change (blue) peaks a few years/generations before selection on species identity (yellow). F) Selection on species rarity (yellow) plus rarity changes (grey) exactly accounts for biodiversity change (blue). G) The frequencies of both species change due to drift (scenario 3). H) In this case, biodiversity (blue) changes little because both species have similar rarity values. As a result, drift’s effects (yellow) on the frequency of species 1 are far stronger than biodiversity change (blue). I) Biodiversity change (blue) is, however, exactly equal to the combined effects of drift (yellow) and rarity changes (yellow).

In open communities, total biodiversity change can be parsed into mechanisms that act on rarity. In a community with immigration (scenario 4), immigrants disproportionately belong to species 1, such that the frequency of species 1 increases despite selection that favors species 2 (Figure 5 A). Over the course of the simulation, biodiversity declines, though this decline slows over time (Figure 5 B). This overall change in biodiversity can be decomposed into a positive effect of immigration. This effect results from the arrival of many individuals of species 1, which is locally rare (Figure 5 C; red dashed line). This effect is offset by selection (Figure 5 C; yellow line) and rarity changes (Figure 5 C; grey line), which decrease biodiversity. In our speciation simulation (scenario 5), species frequencies are more dynamic (Figure 5 D). Over time, change in biodiversity fluctuates around 0 (Figure 5 E). This change roughly parallels drift on species rarity (Figure 5 F; yellow). Speciation increases biodiversity (Figure 5 F; brown dashed line), an effect that is balanced by rarity changes (Figure 5 F; grey line).

**Figure 5.**
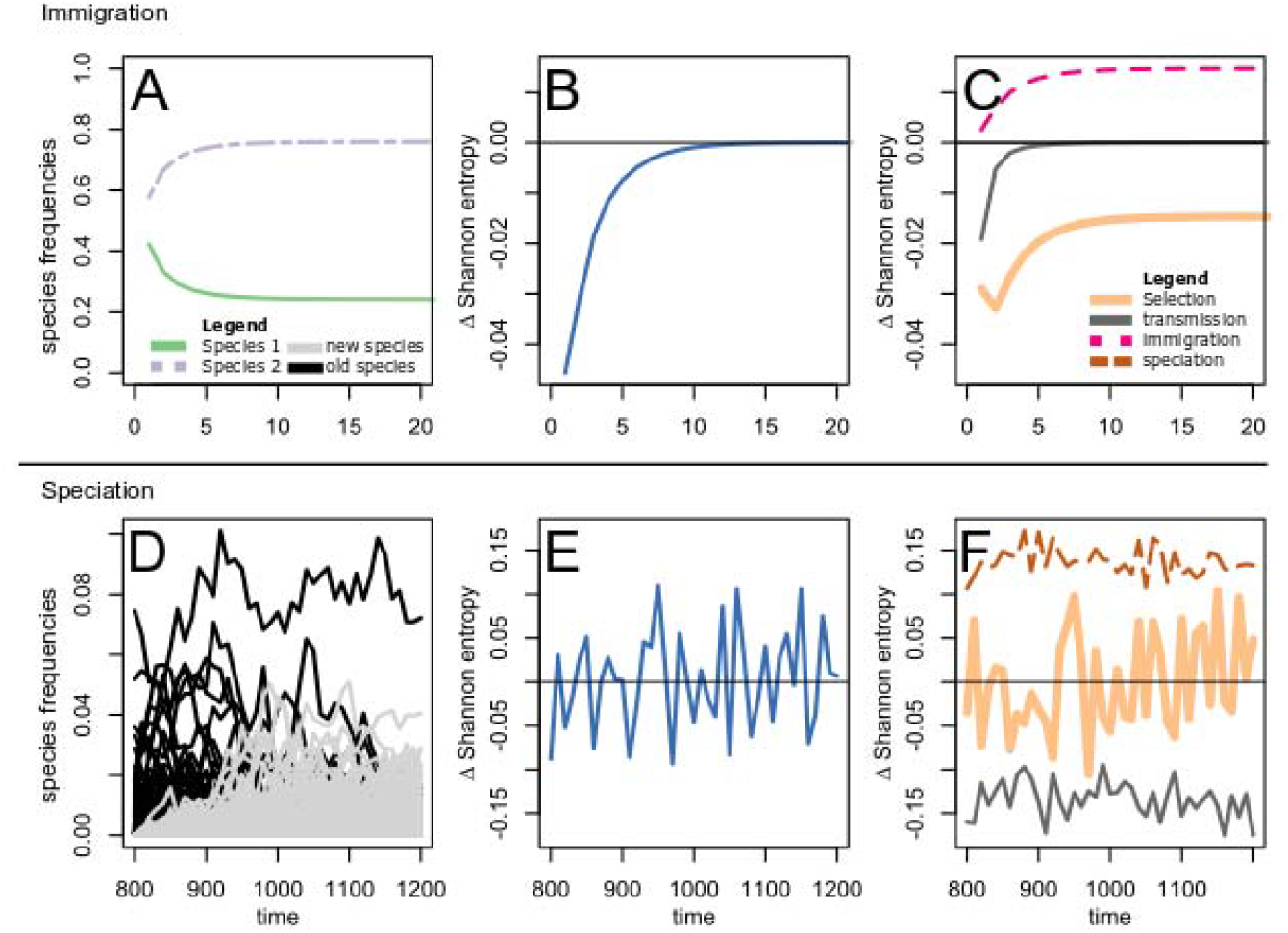
Change in Shannon-Wiener diversity can be decomposed into the effects of multiple mechanisms on rarity. A) The frequency of two species in a single habitat patch subject to selection for species 2 and against species 1 (green). In this patch immigration favours species 1. B) Over the course of the simulation biodiversity (blue) declines. C) This decline can be decomposed into immigration of individuals of the rarer species into the community (pink dashed line) counterbalanced by selection (yellow thick line) and rarity changes (grey line), all of which were measured using Equation 6. D) Frequency change for a community of species subject to drift and speciation. In this panel black lines represent species that were present initially (i.e. when time=800). Grey lines represent new species. E) In this case, total biodiversity change (blue) fluctuates around zero. F) Drift (yellow thick line) roughly mirrors this pattern with an increase in biodiversity due to speciation (brown dashed line), which is counterbalanced by a decrease in biodiversity due to rarity changes (grey solid line), all of which were measured using Equation 6.

## Discussion

Traditional estimates of biodiversity change are low, yet anthropogenic environmental changes seem to pose major threats to biodiversity (Tittensor et al. 2014). This disconnect has generated interest in documenting the mechanisms that shape biodiversity change (Chase and Knight 2013; Hillebrand et al. 2018). Using tools from evolutionary theory, we demonstrate that biodiversity changes in response to mechanisms that act on species’ rarity, and that there is no simple relationship between biodiversity change mechanisms that have traditionally interested ecologists such as competition or selection on species identity. This suggests that biodiversity change can be better understood by quantifying the consequences of mechanisms acting on species rarity. We illustrate this can be done using experiments (Figure 2), field surveys (Figure 3), and simulations (Figure 4, Figure 5).

A major implication of our finding is that changes in rarity often obscure selection’s effects on diversity. Species richness is particularly prone to this problem, with rarity changes counterbalancing extremely strong selection (Figure 1). This result is consistent with previous observations that large changes in species frequencies (for example, due to species’ turnover) can lead to surprisingly small changes in species’ richness (Hillebrand et al. 2018). We show that this problem is more general than previously thought, by showing examples where large changes in species’ frequencies lead to no change in diversity indices. We also illustrate an empirical example of this problem (Figure 3) where strong selection on identity leads to negligible biodiversity change. We then illustrate how our analysis resolves this gap by distinguishing the opposing effects of selection on rarity and rarity changes.

While our approach makes substantial progress towards partitioning sources of biodiversity change, we are unable to distinguish between selection and drift fully. Both processes can produce the same number of descendants for each ancestor (Figure 1 A), which are the relevant quantities tracked by the Price equation. As a result, in our framework, the contributions of drift and selection are considered jointly because additional information is needed to disentangle their effects (Rice 2004). Pure drift occurs when variation in the number of descendants is due to entirely random variation in the number of descendants produced by each type. In contrast, pure selection occurs when all variation in the number of descendants is explained by rarity. In ecology there is no guarantee that selection acts entirely in the absence of drift or vice versa. As with existing analyses of community data, null models can be used to test if the observed patterns are consistent with pure drift (Hillebrand et al. 2018).

Change in the frequency of different types of organisms is fundamental to how selection is formally defined in evolutionary theory (Frank 2012b; Queller 2017; Rice 2004). As a result, formal links between diversity and selection require a way to separate frequency from other facets of diversity. Fortunately, diversity indices cleave into two components: the frequency of individuals belonging to a species, and each individual’s rarity score, as formalized by (Shannon 1948), and described in (Hill 1973; Jost 2007; Patil and Taillie 1982). Though this split is fundamental to how diversity indices have been derived and interpreted, it is counterintuitive because an individual’s rarity score is a function of species frequency.

Our presentation emphasizes the (mathematically valid) distinction between frequency and rarity. This distinction is admittedly subtle because frequency is a function of rarity. For readers who are sceptical of this distinction our results will be of interest because they suggest that diversity change cannot be formally analysed using selection. In evolutionary theory, selection separates frequency change from other sources of change (Frank 2012b). Diversity indexes are functions of frequency and no other variable. If a species’ contribution to diversity cannot be separated into frequency and other components, then the notion of selection cannot be applied rigorously to diversity indexes. Moreover, our work shows that the split between frequency and rarity leads to a mathematically exact partitioning of diversity change, such that the difference in diversity between two sampling periods is exactly equal to the contributions of each of the mechanisms in Equation 6. As such, any method leading to a different answer is likely to lead to paradoxes such as the cases we illustrate (see Figure 1), where selection on individual identity leads to no diversity change, or inaccuracies (see Figure 4 B), where diversity change peaks before selection on individual identity.

For readers willing to accept the distinction between frequency and rarity our approach leads to new tools to study diversity change. We have used Figure 1 to illustrate how our approach can lead to a better understanding of when selection will increase diversity and when it will not. We have shown why selection on species identity is a misleading guide to diversity change in some circumstances (Figure 2,3). This was possible because with our approach, the effects of selection can be quantified from empirical data as can other mechanisms shaping ecological communities. We hope that these advantages will persuade readers to tentatively embrace the split between frequency and rarity.

Selection on rarity captures consequences of many other processes, including both frequency-dependent and frequency-independent selection. This point is emphasized in Figure 4 A-F where two different scenarios are analysed. In both classes total change is partitioned exactly into selection on rarity and rarity change (Figure 4 C, F). This means that once we understand selection on rarity and transmission bias, we have completely described biodiversity change. But in one scenario selection on rarity is a consequence of frequency-independent selection (Figure 4 A) and in the other selection on rarity is and consequence of frequency-dependent selection (Figure 4 D).

Recent works have pointed out a seeming disconnect between mechanisms that shape community ecology and biodiversity change (Vellend et al. 2013, Dornelas et al. 2014, McGill et al. 2015). Existing solutions focus on alternative summaries of change in ecological communities, such as quantifying changes in species turnover between the past and present (Hillebrand et al. 2018). These measures, however, do not clarify how biodiversity itself changes. We believe that more insights can be gained by formally partitioning biodiversity change with the Price equation. In particular, we hypothesize that rarity change frequently opposes the effects of other mechanisms, leading to slower diversity change in nature than would be expected by studying species’ dynamics alone. This effect emerges in several of our examples (Figure 1,3) and is nearly inevitable for the species richness diversity index (Online Appendix S1). These rarity changes are easily missed without a formal partitioning scheme such as the extended Price equation (Equation 6). How frequently rarity changes obscure the effects of other mechanisms in nature remains an open question in ecology. Understanding the importance of this effect will help to clarify when substantial biodiversity change will be observed.

## Supporting information

Supplemental appendix 2 r scripts

## Acknowledgements

Mark Westoby, Thuy Nguyen, and Buck Trible, the other participants in our discussion group, whose thoughts shaped the current article. We are particularly indebted to Margaret Kosmala for creating and organizing these discussions. Our paper would not have been possible without her effort. This manuscript has received thoughtful comments from several previous reviewers, and we are indebted to their work. In particular we recognize Jeremy Fox, who reviewed the manuscript several times, along with James Rosindell and an anonymous reviewer who reviewed the text during an extremely stressful time. Helpful comments were provided by Mark Vellend, Steve Frank, Komei Kadowaki, Erol Ackay, Adrian Paterson, Bevin Brett and Robert Cruickshank. Images of rabbit and squirrel based on illustrations from by Anthony Caravaggi, https://creativecommons.org/licenses/by-nc-sa/3.0/. All illustrations from phylopic http://phylopic.org/.

## Literature cited

Abrams PA (1987) On classifying interactions between populations Oecologia 73:272--281

Adler PB, HilleRisLambers J, Levine JM (2007) A niche for neutrality Ecology Letters 10:95–104

Ayala FJ, Campbell CA (1974) Frequency-dependent selection Annual review of Ecology and systematics 5:115–138

Barton NH, Servedio M (2015) The interpretation of selection coefficients Evolution 69:1101–1112

Blanchet FG, Cazelles K, Gravel D (2020) Co-occurrence is not evidence of ecological interactions Ecology Letters

Chase JM, Knight TM (2013) Scale-dependent effect sizes of ecological drivers on biodiversity: why standardised sampling is not enough Ecology letters 16:17–26

Chase JM, Leibold MA (2003) Ecological niches - linking classical and contemporary approaches. The University of Chicago Press, Chicago IL

Collins S, Gardner A (2009) Integrating physiological, ecological and evolutionary change: a Price equation approach Ecology letters 12:744–757

Conner JK, Hartl DL (2004) A primer of ecological genetics. Sinauer Associates Incorporated,

Cover TM, Thomas JA (2012) Elements of information theory. John Wiley & Sons,

Day T (2005) Modelling the ecological context of evolutionary change: Deja vu or something new? In: Ecological Paradigms Lost: Routes of Theory Change. Elsevier, p 273

Dornelas M, Gotelli NJ, McGill B, Shimadzu H, Moyes F, Sievers C, Magurran AE (2014) Assemblage time series reveal biodiversity change but not systematic loss Science 344:296–299

Engen S, Sæther BE (2014) Evolution in fluctuating environments: decomposing selection into additive components of the Robertson–Price equation Evolution 68:854–865

Frank SA (2012a) Natural selection. III. Selection versus transmission and the levels of selection Journal of evolutionary biology 25:227–243

Frank SA (2012b) Natural selection. IV. The Price equation Journal of evolutionary biology 25:1002–1019

Frank SA (2012c) Natural selection. V. How to read the fundamental equations of evolutionary change in terms of information theory Journal of evolutionary biology 25:2377–2396

Genung MA, Schweitzer JA, Ubeda F, Fitzpatrick BM, Pregitzer CC, Felker-Quinn E, Bailey JK (2011) Genetic variation and community change–selection, evolution, and feedbacks Functional Ecology 25:408–419

Govaert L, Pantel JH, De Meester L (2016) Eco-evolutionary partitioning metrics: assessing the importance of ecological and evolutionary contributions to population and community change Ecology letters 19:839–853

Hedrick PW (2011) Genetics of populations. Jones & Bartlett Learning,

Hill MO (1973) Diversity and evenness: a unifying notation and its consequences Ecology 54:427–432

Hillebrand H et al. (2018) Biodiversity change is uncoupled from species richness trends: Consequences for conservation and monitoring Journal of Applied Ecology 55:169–184

Hubbell SP (2001) A unified neutral theory of biodiversity and biogeography. Princeton University Press, Princeton, NJ

Jablonski D (2008) Species selection: theory and data Annual review of ecology, evolution, and systematics 39:501–524

Jia X, Dini-Andreote F, Salles JF (2018) Community assembly processes of the microbial rare biosphere Trends in microbiology 26:738–747

Johnson MTJ, Stinchcombe JR (2007) An emerging synthesis between community ecology and evolutionary biology Trends in Ecology and Evolution 22:250–257

Jost L (2006) Entropy and diversity Oikos 113:363–375

Jost L (2007) Partitioning diversity into independent alpha and beta components Ecology 88:2427–2439

Kahn_Academy_Labs (2014) Information entropy | Journey into information theory | Kahn Academy.

Kerr B, Godfrey-Smith P (2009) Generalization of the Price equation for evolutionary change Evolution 63:531–536

Lande R (1979) Quantitative genetic analysis of multivariate evolution, applied to brain: body size allometry Evolution 33:402–416

Lande R, Arnold SJ (1983) The measurement of selection on correlated characters Evolution 37:1210–1226

Lehtonen J (2018) The Price Equation, Gradient Dynamics, and Continuous Trait Game Theory The American Naturalist 191:146–153

Loreau M, Hector A (2001) Partitioning selection and complementarity in biodiversity experiments Nature 412:72–76

Luís R, Elaydi S, Oliveira H (2011) Stability of a Ricker-type competition model and the competitive exclusion principle Journal of Biological Dynamics 5:636–660

MacArthur R (1970) Species packing and competitive equilibrium for many species Theoretical population biology 1:1–11

MacArthur RH (1972) Geographical Ecology: Patterns in the Distribution of Species. Harper & Row, New York

MacDonald A, Vellend M (2016) R code for Chapter 6: Simulating Dynamics in Ecological Communities. Accessed 11 November 2017 2017

Magurran AE (2013) Measuring biological diversity. John Wiley & Sons,

Mallet J (2012) The struggle for existence: how the notion of carrying capacity, K, obscures the links between demography, Darwinian evolution, and speciation Evolutionary Ecology Research 14:627–665

McGill BJ, Dornelas M, Gotelli NJ, Magurran AE (2015) Fifteen forms of biodiversity trend in the Anthropocene Trends in ecology & evolution 30:104–113

Norberg J, Urban MC, Vellend M, Klausmeier CA, Loeuille N (2012) Eco-evolutionary responses of biodiversity to climate change Nature Climate Change 2:747–751

Nowak MA (2006) Evolutionary dynamics. Harvard University Press,

Otto SP, Day T (2007) A biologist’s guide to mathematical modeling in ecology and evolution vol 13. Princeton University Press,

Patil G, Taillie C (1982) Diversity as a concept and its measurement Journal of the American statistical Association 77:548–561

Perring MP et al. (2016) Global environmental change effects on ecosystems: the importance of land-use legacies Global change biology 22:1361–1371

Price GR (1970) Selection and covariance Nature 227:520–521

Price GR (1972) Extension of covariance selection mathematics Annals of human genetics 35:485–490

Price GR (1995) The nature of selection Journal of Theoretical Biology 175:389–396

Purves DW, Turnbull LA (2010) Different but equal: the implausible assumption at the heart of neutral theory Journal of Animal Ecology 79:1215–1225

Queller DC (2017) Fundamental Theorems of Evolution The American Naturalist 189:345–353

R Core Team (2019) R: A Language and Environment for Statistical Computing.

Rabosky DL, McCune AR (2010) Reinventing species selection with molecular phylogenies Trends in Ecology & Evolution 25:68–74

Rankin BD et al. (2015) The extended Price equation quantifies species selection on mammalian body size across the Palaeocene/Eocene Thermal Maximum Proc R Soc B 282:20151097

Remus-Emsermann MN, Pelludat C, Gisler P, Drissner D (2018) Conjugation dynamics of self-transmissible and mobilisable plasmids into <em>E. coli</em> 0157:H7 on <em>Arabidopsis thaliana</em> rosettes bioRxiv doi:10.1101/375402

Rice SH (2004) Evolutionary theory: mathematical and conceptual foundations. Sinauer Associates Sunderland,

Ricker WE (1954) Stock and recruitment Journal of the Fisheries Board of Canada 11:559–623

Rosindell J, Hubbell SP, Etienne RS (2011) The unified neutral theory of biodiversity and biogeography at age ten Trends in ecology & evolution 26:340–348

Rosindell J, Hubbell SP, He F, Harmon LJ, Etienne RS (2012) The case for ecological neutral theory Trends in ecology & evolution 27:203–208

Shannon CE (1948) A mathematical theory of communication Bell system technical journal 27:379–423

Sherwin WB, Chao A, Jost L, Smouse PE (2017) Information theory broadens the spectrum of molecular ecology and evolution Trends in ecology & evolution 32:948–963

Sirianni K (2018) Connecting zooplankton populations in time and space: How dispersal, biotic, and abiotic factors influence metapopulation dynamics Tilman D (1980) A graphical-mechanistic approach to competition and predation The American Naturalist 116:362–393

Tilman D, Wedin D (1991) Dynamics of nitrogen competition between successional grasses Ecology 72:1038–1049

Tittensor DP et al. (2014) A mid-term analysis of progress toward international biodiversity targets Science 346:241–244

Traulsen A, Nowak MA (2006) Evolution of cooperation by multilevel selection Proceedings of the National Academy of Sciences 103:10952–10955

Vellend M (2010) Conceptual synthesis in community ecology The Quarterly review of biology 85:183–206

Vellend M (2016) The theory of ecological communities (MPB-57). Princeton University Press,

Vellend M et al. (2017) Plant biodiversity change across scales during the Anthropocene Annual Review of Plant Biology 68

Vellend M et al. (2013) Global meta-analysis reveals no net change in local-scale plant biodiversity over time Proceedings of the National Academy of Sciences 110:19456–19459

