## Supplemental appendix 1 for "Selection and Biodiversity change"

### Appendix S1

#### S1.1 Selection leaves species richness unchanged.

The most common measures of species diversity can be expressed as measurements of average rarity. As such, they are amenable to analyses with the Price equation. To express species richness in the format we get: $z_{i}=\frac{1}{p_{i}}$ such that:

$$\sum_{i} p_{i}\frac{1}{p_{i}}$$

$$=\sum_{i} 1$$

$$=i$$

Where *i* is the number of species. In the case that all species survive to the present (i.e. $p_{i}^{'}>0$), we can compute change as:

(1) $\overset{Change}{\overbrace{\Delta\left( \sum_{i} p_{i}z_{i} \right)}}=\overset{Selection}{\overbrace{\sum_{i} \Delta p_{i}z_{i}}}+\overset{Transmission}{\overbrace{\sum_{i} p_{i}'\Delta z_{i}}}$

When we substitute $z_{i}=\frac{1}{p_{i}}$

$$\sum_{i} \Delta p_{i}\frac{1}{p_{i}}+\sum_{i} p_{i}'\left( \frac{1}{p_{i}'}-\frac{1}{p_{i}} \right)$$

$$=\sum_{i} \left( \frac{p_{i}'}{p_{i}}-\frac{p_{i}}{p_{i}} \right)+\sum_{i} \left( \frac{p_{i}'}{p_{i}'}-\frac{p_{i}'}{p_{i}} \right)$$

$$=\sum_{i} \left( \frac{p_{i}'}{p_{i}}-1 \right)+\sum_{i} \left( 1-\frac{p_{i}'}{p_{i}} \right)$$

=0

This demonstrates that the selection term and the transmission term cancel out exactly when studying species richness, as long as all species are present in the descendant species pool.

#### S1.2 Selection can leave all diversity indices unchanged

This example considers a community of two species which exchange rarity values due to selection on species identity. In the past: $p_{1}=x$ and $p_{2}=y$. In the present: $p_{1}'=y$ and $p_{2}'=x$. When the frequency of a species is *x* we define its rarity score to be a function of that frequency: *f(x)*. Different functions will lead to different diversity indices. For example : *f(x)=x* for Simpson’s diversity and *f(x)=-log(x)* for Shannon Wiener.

Substituting these definitions into Equation S1 gives:

(2) $\underset{selection}{\underbrace{\left( y-x \right)f\left( x \right)+\left( x-y \right)f\left( y \right)}}+\underset{transmission}{\underbrace{y\left( f\left( y \right)-f\left( x \right) \right)+x\left( f\left( x \right)-f\left( y \right) \right)}}$

Expanding we obtain:

(3) $\underset{selection}{\underbrace{yf\left( x \right)-xf\left( x \right)+xf\left( y \right)-yf\left( y \right)}}+\underset{transmission}{\underbrace{-yf\left( x \right)+yf\left( y \right)-xf\left( y \right)+xf\left( x \right)}}$

These two terms cancel out exactly, which means that total change in diversity is 0 regardless of which function is used to measure rarity.

#### S1.3 Strength of selection on species identity

Equation 3 can also be used to quantify selection on species identity for a species of interest. To do this label the species of interest species 1. Then, let $z_{i}$ be an indicator variable taking on a value of 1 when *i* =1 and 0 otherwise (Nowak 2006, Traulsen and Nowak 2006). In the absence of speciation, species identity remains fixed such that $\Delta z_{i}=0$. Substituting these terms into Equation S1 from the main text we find that:

(4) $\overset{Change}{\overbrace{\Delta\left( \sum_{i} p_{i}z_{i} \right)}}=\overset{Selection}{\overbrace{\Delta p_{1}}}$

Note that selection on identity implies a separate measurement of selection for each species in a community, or more precisely n-1 measurements in a community of n species. Once the strength of selection on n-1 species is known, the strength of selection on the final species can be deduced. In contrast, selection on rarity provides a single summary of biodiversity change across the entire community.

Nowak, M. A. 2006. Evolutionary dynamics. Harvard University Press.

Traulsen, A., and M. A. Nowak. 2006. Evolution of cooperation by multilevel selection. Proceedings of the National Academy of Sciences **103**:10952-10955.
