## Supplemental appendix 2 r scripts for "Selection and Biodiversity change": Compare selection on identity rarity.Rmd

````
---
title: "Godsoe et al. the fifth mechanism"
author: "William Godsoe"
output:
  html_document: default
  pdf_document: default
  word_document: default
---

```{r setup, include=FALSE}
knitr::opts_chunk$set(echo = TRUE,fig.path = "figExamples/")
require(RColorBrewer)
require(knitr)
#
#requires: RColorBrewer, knitr
#input: "Example 1.R", "Example 5.R", "Example 7.R", 
"functionsforVspeciationeg.R"

#This file is used to call scripts for each of the simulations presented in the main text and supplements of Godsoe et al. submitted.
#once the simulations have run, this script processeses and plots the results. 
#you will need to change the working directory to the appropriate location using a setwd() command

#a reminder that it may be desirable to change the output format from pdf to html depending on the software on your computer. 
#this can be done (in Rstudio) using the "Knit" button


#you should set the working directory to the spot where all the scripts from this project are stored. This can be done
#using a setwd() command. Below i've indicated where to do this


```


```{r eg1,fig.height = 4, fig.width = 6,warning=FALSE, echo=FALSE} 
#, results='hide',echo=F


rm(list=ls())


######################################################################################
###################################################################################################Here is where you must set the appropriate directory ####################
######################################################################################
######################################################################################
######################################################################################

#setwd("")
source("functionsforVspeciationeg.R")

rcol<-brewer.pal(8, "Accent")

#these parameters control the scope of the simulation (i.e. number of generations, patches etc.)
num.patch <- 1
generations<-50
num.years<-generations+1
times<-1:(generations+1)
pDispersToPatch1<-NA


#these parameters are used to describe population growth, and are explained in the main text
r1=0.3
r2=0.3
a11=0.001 
a12=0.001
a21=0.001
a22=0.001


#the following script simulates population growth and calculates terms in the price equation
source("Example 1 equal ish ini.R")


#the following are notes on teh three ways we can use to compute teh change in numbers equivalent of shannon weiner
#diversity:


#shannonRatio<-1:50
#for(i in 1:50){
#  shannonRatio[i]<-exp(shannonDiversity[i+1])/exp(shannonDiversity[i])
#}

drift<-list(pNi=pNi,
            partitionSimpsons=partitionSimpsons,
            simpsonssChange4mech=simpsonssChange4mech,
            simpsonsChange=simpsonsChange,
            partitionSW=partitionSW,
            shannonChange=shannonChange,
            shannonChange4mech=shannonChange4mech,
            indBiomass=indBiomass,
            partitionBiomass=partitionBiomass,
            biomassChange=biomassChange,
            biomassChange4mech=biomassChange4mech)


#the following are notes on teh three ways we can use to compute teh change in numbers equivalent of shannon weiner
#diversity:


#shannonRatio<-1:50
#for(i in 1:50){
#  shannonRatio[i]<-exp(shannonDiversity[i+1])/exp(shannonDiversity[i])
#}


source("Example 1 equal ish ini.R")

drift1<-list(pNi=pNi,
            partitionSimpsons=partitionSimpsons,
            simpsonssChange4mech=simpsonssChange4mech,
            simpsonsChange=simpsonsChange,
            partitionSW=partitionSW,
            shannonChange=shannonChange,
            shannonChange4mech=shannonChange4mech,
            indBiomass=indBiomass,
            partitionBiomass=partitionBiomass,
            biomassChange=biomassChange,
            biomassChange4mech=biomassChange4mech)


source("Example 1 equal ish ini.R")

drift2<-list(pNi=pNi,
            partitionSimpsons=partitionSimpsons,
            simpsonssChange4mech=simpsonssChange4mech,
            simpsonsChange=simpsonsChange,
            partitionSW=partitionSW,
            shannonChange=shannonChange,
            shannonChange4mech=shannonChange4mech,
            indBiomass=indBiomass,
            partitionBiomass=partitionBiomass,
            biomassChange=biomassChange,
            biomassChange4mech=biomassChange4mech)


num.patch <- 1
generations<-50
num.years<-generations+1
times<-1:(generations+1)
pDispersToPatch1<-NA

#to represent frequency independent selection set r1>r2 indicating higher fitness of species 1
r1=0.6
r2=0.3

a11=0.001
a12=0.001
a21=0.001
a22=0.001


source("Example 1.R")

selectionFI<-list(pNi=pNi,
            partitionSimpsons=partitionSimpsons,
            simpsonssChange4mech=simpsonssChange4mech,
            simpsonsChange=simpsonsChange,
            partitionSW=partitionSW,
            shannonChange=shannonChange,
            shannonChange4mech=shannonChange4mech,
            indBiomass=indBiomass,
            partitionBiomass=partitionBiomass,
            biomassChange=biomassChange,
            biomassChange4mech=biomassChange4mech)

source("Example 5.R")

immigration<-list(pNi=pNi,
            partitionSimpsons=partitionSimpsons,
            simpsonssChange4mech=simpsonssChange4mech,
            simpsonsChange=simpsonsChange,
            partitionSW=partitionSW,
            shannonChange=shannonChange,
            shannonChange4mech=shannonChange4mech,
            indBiomass=indBiomass,
            partitionBiomass=partitionBiomass,
            biomassChange=biomassChange,
            biomassChange4mech=biomassChange4mech)


num.patch <- 1
generations<-50
num.years<-generations+1
times<-1:(generations+1)
pDispersToPatch1<-NA


r1=0.3
r2=0.3

#note that interspecific competition is weaker than intraspecific competition
a11=0.001
a12=0.0001
a21=0.0001
a22=0.001


source("Example 1.R")


selectionFD<-list(pNi=pNi,
            partitionSimpsons=partitionSimpsons,
            simpsonssChange4mech=simpsonssChange4mech,
            simpsonsChange=simpsonsChange,
            partitionSW=partitionSW,
            shannonChange=shannonChange,
            shannonChange4mech=shannonChange4mech,
            indBiomass=indBiomass,
            partitionBiomass=partitionBiomass,
            biomassChange=biomassChange,
            biomassChange4mech=biomassChange4mech)


```


```{r diversity,echo=FALSE, fig.height=6,fig.width=8,eval=TRUE, cache=F}

#setup for including numbers equivalent height 7 width 10
#par(mfrow=c(5,6), mar=c(3, 3.7, 0, 0) + 0.1)

par(mfrow=c(3,3), mar=c(5, 4.5, 0, 0) + 0.1)

diversityRow<-function(simData,
                    mains=NA,
                    biomassCol=rcol[1:2],
                    deltaCol=rcol[c(3,NA,NA,5,NA)],
                    xtext="",
                    mech4lab=NA,
                    ylabBottom=NA,
                    ylaC2=NA,
                    y1=NA,
                    partition=NA){
  
  
  #plot change in frequency of two species
  
  plot(simData$pNi[,1], type="l", ylim=c(0,1), ylab="", xlab=xtext, col=rcol[1],lwd=2, lty=1)
  points(simData$pNi[,2], type="l",  xlab="time", col=rcol[2],lwd=2, lty=6)
  title(ylab=y1,line=2)

  #partition$total<-rowSums(partition)
  
  
  yrange<-range(c(range(partition),range(diff(simData$pNi[,1]))))

  plot(diff(simData$pNi[,1]), type="l", ylab=ylaC2, xlab=xtext, col=deltaCol[1],lwd=3, lty=1,ylim=yrange)
  points(partition$divChange, type="l", ylab="", xlab="", col=deltaCol[4],lwd=3, lty=1)

  abline(h=0)
 # points(simData$pNi[,2], type="l",  xlab="time", col=rcol[2],lwd=2, lty=6)
 # title(ylab=y1,line=2)


#plot overall diviersity change. 
  matplot(1:generations, partition, type = "l", 
        xlab = xtext, 
        ylab = ylabBottom, lty=1,col=deltaCol,lwd=c(3,0,0,3))
  abline(h=0, col="black", lwd=1)
  
LS<-length(partition[,1])
  
selPTran<-partition[,1]+partition[,3]
abline(h=0)


polygon(c(1:LS,LS:1),
        c(partition[,1],selPTran[LS:1]),
        col="lightgrey",
        border=NA)


  
 # plot(diff(simData$pNi[,1]), simData$partitionSW$total,  ylab="", xlab="", col="darkgrey",lwd=2, lty=1, pch=16)
 # points(simData$partitionSW[,1], simData$partitionSW$total,  ylab="", xlab="", col="blue",lwd=2, lty=1, pch=16)

#abline(h=0)
#abline(0,1)
  
  return()
  }

#svg("SelectionDrift.svg", width=6, height=6)

#diversityRow(selectionFI,y1="1) selection \n(freq. independent)")
#diversityRow(selectionFD,y1="2) selection \n(freq. dependent)")
#diversityRow(drift,y1="3) drift", xtext="Time",ylaC2="selection on identity",
#            ylabBottom = expression(paste( Delta," Shannon entropy")))
diversityRow(selectionFI,y1="1) selection \n(freq. independent)", partition=selectionFI$partitionSW)
diversityRow(selectionFD,y1="2) selection \n(freq. dependent)", partition=selectionFD$partitionSW)
diversityRow(drift,y1="3) drift", xtext="Time",ylaC2="selection on identity",
             ylabBottom = expression(paste( Delta," Shannon entropy")), partition=drift$partitionSW)

#

 # dev.off()

  
#svg("SelectionDriftSimpsons.svg", width=6, height=6)

par(mfrow=c(3,3), mar=c(5, 4.5, 0, 0) + 0.1)

  
diversityRow(selectionFI,y1="1) selection \n(freq. independent)", partition=selectionFI$partitionSimpsons)
diversityRow(selectionFD,y1="2) selection \n(freq. dependent)", partition=selectionFD$partitionSimpsons)
diversityRow(drift,y1="3) drift", xtext="Time",ylaC2="selection on identity",
             ylabBottom = expression(paste( Delta," Simpsons")), partition=drift$partitionSimpsons)
#dev.off()

```


```{r diversity1,echo=FALSE, fig.height=3,fig.width=6, cache=T}
#################################################################################
#svg("New arrivals.svg", width=6, height=4)
par(mfrow=c(2,3), mar=c(3, 3.7, 0, 0) + 0.1)


diversityRow<-function(simData,
                    mains=NA,
                    biomassCol=rcol[1:2],
                    deltaCol=rcol[c(3,NA,8,5,NA)],
                    xtext=NA,
                    mech4lab=NA,
                    ylabBottom=NA,
                    y1=NA,
                    partition=NA){
  
  
  #plot change in frequency of two species
  
  plot(simData$pNi[,1], type="l", ylim=c(0,1), ylab="", xlab="", col=rcol[1],lwd=2, lty=1, xlim=c(0,20))
  points(simData$pNi[,2], type="l",  xlab="time", col=rcol[2],lwd=2, lty=6)
  title(ylab=y1,line=2)

  partition$total<-rowSums(partition)
  
  
  yrange<-range(c(range(partition$total),range(diff(simData$pNi[,1]))), xlim=c(0,20))

 # plot(diff(simData$pNi[,1]), type="l", ylab="", xlab="", col=deltaCol[1],lwd=4, lty=1,ylim=yrange)
 # points(simData$partitionSW$total, type="l", ylab="", xlab="", col=deltaCol[4],lwd=2, lty=1)

#  abline(h=0)
 # points(simData$pNi[,2], type="l",  xlab="time", col=rcol[2],lwd=2, lty=6)
 # title(ylab=y1,line=2)


#plot overall diviersity change. 
  matplot(1:generations, partition, type = "l",  xlim=c(0,20),
        xlab = "", 
        ylab = "", lty=1,col=rcol[c(NA,NA,NA,5,NA)],lwd=c(2,2,2,2))
  abline(h=0, col="black", lwd=1)
  
#plot overall diviersity change. 
  matplot(1:generations, partition, type = "l", xlim=c(0,20),
        xlab = "", 
        ylab = "", lty=c(1,2,1,1,1),col=rcol[c(3,6,8,NA,NA)],lwd=c(4,2,2,2))
  abline(h=0, col="black", lwd=1)
  
  
 # plot(diff(simData$pNi[,1]), simData$partitionSW$total,  ylab="", xlab="", col="darkgrey",lwd=2, lty=1, pch=16)
 # points(simData$partitionSW[,1], simData$partitionSW$total,  ylab="", xlab="", col="blue",lwd=2, lty=1, pch=16)

abline(h=0)
#abline(0,1)
  
  return()
  }


diversityRow(immigration,deltaCol=rcol[c(3,6,8,5,NA)],y1="4) immigration",partition=immigration$partitionSimpson)

source("Example 7.R")


tRange<-80:120
#black species were in the community at the start of the time period tRange,
#red species were not, and so could be new
possibleNewSpecies<-ifelse(pNi[min(tRange),]!=0, "black","lightgrey")
possibleNewSpecies[colSums(pNi[tRange,])==0]<-NA #ignore species not present durign tRange

yearRange<-tRange*sampFreq
plotTime<-20
partitionSW$total<-rowSums(partitionSW)
partitionSimpsons$total<-rowSums(partitionSimpsons)


matplot(yearRange, pNi[tRange,], type = "l", 
        xlab = "Time", 
        ylab = "", lty=1,col=possibleNewSpecies,lwd=2, ylim=c(0,max(pNi[tRange,])))
  title(xlab="time",ylab=expression(paste("5) speciation")), line=2, cex.lab=1)


  matplot(yearRange, partitionSW[tRange,], type = "l", 
        xlab = "", 
        ylab = "" ,lty=1,
        col=rcol[c(NA,NA,NA,5,NA)],
        #col=c(rcol[3],"lightgrey",rcol[8]),
        lwd=c(2,1,1,1)*2)
abline(h=0, col="black", lwd=1)
  title(xlab="time", ylab=expression(paste( Delta," Shannon entropy")), line=2, cex.lab=1)
  
matplot(yearRange, partitionSW[tRange,], type = "l", 
        xlab = "", 
        ylab = "" ,lty=c(1,5,1,1,1),
        col=rcol[c(3,7,8,NA,NA)],
        #col=c(rcol[3],"lightgrey",rcol[8]),
        lwd=c(2,1,1,2)*2)
abline(h=0, col="black", lwd=1)
  title(xlab="time", ylab=expression(paste( Delta," Shannon entropy")), line=2, cex.lab=1)


  
  
#########plot of simpsons diversity###################
  diversityRow(immigration,deltaCol=rcol[c(3,6,8,5,NA)],y1="4) immigration",partition=immigration$partitionSW)

  
  
  
  
  matplot(yearRange, pNi[tRange,], type = "l", 
        xlab = "Time", 
        ylab = "", lty=1,col=possibleNewSpecies,lwd=2, ylim=c(0,max(pNi[tRange,])))
  title(xlab="time",ylab=expression(paste("5) speciation")), line=2, cex.lab=1)


  matplot(yearRange, partitionSimpsons[tRange,], type = "l", 
        xlab = "", 
        ylab = "" ,lty=1,
        col=rcol[c(NA,NA,NA,5,NA)],
        #col=c(rcol[3],"lightgrey",rcol[8]),
        lwd=c(2,1,1,1)*2)
abline(h=0, col="black", lwd=1)
  title(xlab="time", ylab=expression(paste( Delta," Shannon entropy")), line=2, cex.lab=1)
  
matplot(yearRange, partitionSimpsons[tRange,], type = "l", 
        xlab = "", 
        ylab = "" ,lty=c(1,5,1,1,1),
        col=rcol[c(3,7,8,NA,NA)],
        #col=c(rcol[3],"lightgrey",rcol[8]),
        lwd=c(2,1,1,2)*2)
abline(h=0, col="black", lwd=1)
  title(xlab="time", ylab=expression(paste( Delta," Simpsons")), line=2, cex.lab=1)

  
  
  
  
  
 # dev.off()
```
````
