## Supplemental appendix 2 r scripts for "Selection and Biodiversity change": functionsforVspeciationeg.R

```
####function declarations######

#this function modifies R's default covariance in two ways
#first it calculates populatoin covariance, not sample covariance
#second it checks to see if the legth of the datasets is 1, by default
#r returns NA in this case, but mathematically the covariance should be 0
#since x=xbar and y=ybar so the standard cov formula becomes 0.
cov.pop <- function(x,y=NULL) {
  
  if(length(x)>1){
    cValue<-cov(x,y)*(length(x)-1)/length(x)
  }
  else{
    cValue<-0
  }
  
  return(cValue)
}


findACOnnections <- function(iTag,COMstart1) {
  #compile a nicely formatted dataframe of the # of decendant individuals connected with each
  #ancestor
  aConnections<-as.data.frame(table(iTag)) 
  aConnections$iTag<-as.numeric(aConnections$iTag)
  
  #find the species associated with each ancestral individual
  aConnections$sA<-as.factor(COMstart1[aConnections$iTag])
  names(aConnections)<-c("individual","con","species")                       
  
  
  #compute the frequency of each species at the start of the study
  pA<-table(COMstart1)/J
  
  aConnections$Pi<-pA[aConnections$species]
  #look up the frequency of each species in the table
  #aConnections$Pi<-pA[aConnections$species]
  
  return(aConnections)
}


#this function recieves a tag for each individual and the abundance of each species in the decendant population
#
#it produces a table including every individual in the decendant community
#It then counts how many connections associate each decendant
#with ancestors. Effectively this is 1 if the species
#was in teh community before and 0 otherwise.
#I note the species associated with each individual. 
#One can then add traits into each individual. As of June 2017, I add the frequency of an individual's species


findDConnections <- function(iTag,COMstart1) {
  dConnections<-data.frame(indTag=1:J,
                           Freq=as.numeric(!is.na(iTag)), #this line checks for new species (NA's) then gives
                           #all other individuals a value of 1.
                           SA=as.factor(COMstart1)) #recheck
  names(dConnections)<-c("individual","con","species")  
  
  pD<-table(COMstart1)/J
  
  dConnections$Pi<-pD[dConnections$species]
  return(dConnections)
}

calculateTraitChange<- function(aCon,dCon) {
  #this funciton is designed to compute "second" term in the denominator of
  #K&G Page 531 equation 1.
  
  #to do this i take summaries of the ancesstral and decendant connections
  
  #create a new vector for aConn which takes each ancestral individual
  #looks up its species then notes frequency of that species in teh decendant
  #community
  aCon$Pidecendant<-dCon[match(aCon$species,dCon$species),'Pi']
  
  #NAs can emerge when a species is absent in the decendant population i.e. if it has gone extinct
  #to fix this i'm adding a 0 here. 
  
  aCon$Pidecendant[is.na(aCon$Pidecendant)]<-0 
  
  #calculate the average deviation in character between ancestral entity i and its connected
  #decendant entities
  #note this formula has a simplification from that presented on P532 of Kerr
  #this should be a correct calculation of (deltaX)^i in Kerr Godfrey smith
  #but only for the special case that all decendants have the same trait values
  #so that sum(cij(Xj-Xi))/ci* reduces to ci*(Xj-Xi)/ci* =(Xj-Xi)
  deltaXi<-(aCon$Pi-aCon$Pidecendant)
  #print((aCon$con)[9132])
  #print((aCon$con*deltaXi)[9132])
  
  term2<-mean(aCon$con*deltaXi)
  return(term2)
  
}


ePrice<-function(cA, cD, traitA,traitD,term2){
  #this function takes the connections associated with each ancestor,
  #the connections associated iwth each decendant, and the traits of
  #ancestral community and decendant community and trait change
  #then resturns each term of the price equation
  #total number of connections associated with ancestors
  cAStar<-sum(cA)
  cStarD<-sum(cD)
  
  cStarStar<-sum(cAStar)
  
  #number of entities
  nA<-length(traitA)
  nD<-length(traitD)
  
  term1<- cov.pop(cA,traitA)
  term3<- -(nD/nA)*cov.pop(cD,traitD)
  return(c(term1,term2,term3)/(cStarStar/nA))
}

SpeciesRichness<-function(U){
  return(length(unique(U)))
}


#functions to add

calculateShannon<-function(Ni){
  Ni<-Ni[Ni>0]
  Ni<- -(Ni)*log(Ni)
  return(sum(Ni))
}


#designed to read line vellends output file and calculate simpsons
calcSimpsons1 <- function(Ni) {
  N1<-sum((table(Ni)/J)^2)
  return(N1)
}


calcSimpsons <- function(Ni) {
  Ni<-(Ni)^2
  return(sum(Ni))
}
classicPrice <- function(qa,qd,za,zd) {
  #since qd=qa(wi/wbar)
  #and wbar should be 1 in a constant population
  
  #relative fitness
  relFitness=qd/qa
  deltaZ<-zd-za
  
  
  #note the computatoin of the price eq is slightly modified to
  #take terms as described in frank 2012 just above eq 2.
  term1<-sum(qa*(relFitness-1)*za)
  term3<-sum(qa*relFitness*(deltaZ))
  
  pTerms<-c(term1,term3)
  return(pTerms)
}
classicPriceExtended <- function(qa,qd,alpha1,za,zd,zStar,P) {
  
  #frequency in ancestral pop
  #frequency of decendants of ancestral pop
  #frequency of immigrants in species j
  #z trait value ancestors
  #trait value of decendants
  #trait value of immigrants
  #proportion of immigrants
  
  zaBar<-sum(qa*za)
  zdBar<-sum(qd*zd)
  
  #first calculate terms involving non-immigrants
  #note qd is corrected so that it only includes entries that match qa
  #as elsewhere my merging is imperfect. 
  wi=qd[1:length(qa)]/qa
  relFitness=qd[1:length(qa)]/qa
  deltaZ<-zd[1:length(qa)]-za
  
  
  term1<-sum(qa*(relFitness-1)*za)*(1-P)
  term3<-sum(qa*wi*(deltaZ))*(1-P)
  
  #convert dimensions to those of final popl
  qa <- c(qa, rep(0, length(alpha1)-length(qa)))
  qd <- c(qd, rep(0, length(alpha1)-length(qd)))
  za <- c(za, rep(0, length(alpha1)-length(za)))
  zd <- c(zd, rep(0, length(alpha1)-length(zd)))
  
  
  #since qd=qa(wi/wbar)
  #and wbar should be 1 in a constant population
  zBar<-sum(qa*za) #average trait value in decendant population
  ZBarSt<-sum(alpha1*zStar)
  
  term2<-P*(ZBarSt-zBar)
  pTerms<-c(term1,term2,term3)
  return(pTerms)
}


classicPriceExtended1 <- function(qa,qd,alpha1,za,zd,zStar,P) {
  
  #frequency in ancestral pop
  #frequency of decendants of ancestral pop
  #frequency of immigrants in species j
  #z trait value ancestors
  #trait value of decendants
  #trait value of immigrants
  #proportion of immigrants

  #first reformat to calculate trait change with "old species" by weeding out species only found among
  #decendants
  qaC<-qa[qa>0]
  qdC<-qd[qa>0]
  zaC<-za[qa>0]
  
  #first calculate terms involving non-immigrants
  wi=qdC/qaC                                  
  deltaZ<-zd-za
  

  #ignore "new species"
  deltaZ<-deltaZ[qa>0]
  
  
  term1<-sum(qaC*(wi-1)*zaC)*(1-P)
  term3<-sum(qaC*wi*(deltaZ))*(1-P)
  
  if(anyNA(alpha1)==FALSE){
  #since qd=qa(wi/wbar)
  #and wbar should be 1 in a constant population
    zBar<-sum(qa*za) #average trait value in ancestral population
    ZBarSt<-sum(alpha1*zStar)
    term2<-P*(ZBarSt-zBar)
  }
  else{
    term2<-0
  }

  pTerms<-c(term1,term2,term3)

  return(pTerms)
}

classicPriceExtendedSW <- function(qa,qd,alpha1,za,zd,zStar,P) {
  
  #frequency in ancestral pop
  #frequency of decendants of ancestral pop
  #frequency of immigrants in species j
  #z trait value ancestors
  #trait value of decendants
  #trait value of immigrants
  #proportion of immigrants
  
  #first reformat to calculate trait change with "old species" by weeding out species only found among
  #decendants
  qaC<-qa[qa>0]
  qdC<-qd[qa>0]
  zaC<- -log(qa[qa>0])
  zdC<- -log(qd[qa>0]) #trait values of decendants of ancestral community
  
  #first calculate terms involving non-immigrants
  # wi=qdC/qaC                                  
  #deltaZ<-zdC-zaC
  deltaQi<-qdC-qaC #change in frequency values form ancestor to decendant
  
  #ignore "new species"
  #deltaZ<-deltaZ[qa>0]
  
  
  term1<-sum(deltaQi*zaC)*(1-P)
  
  #term 2
  
  if(anyNA(alpha1)==FALSE){
    #since qd=qa(wi/wbar)
    #and wbar should be 1 in a constant population
    zBar     <-sum(qaC*zaC) #average trait value in ancestral population
    #the frequency of new species is alpha times the proportion of individuals in new sp in community
    zStari   <- -log(alpha1[alpha1>0]*P)
    zStarBar <-sum(alpha1[alpha1>0]*zStari)
    
    # print(zStari)
    term2    <-(zStarBar -zBar)*P
  }
  else{
    term2<-0
  }
  
  
  #here i am calculating sum(q'(z'-z)) which for Shannon weiner is sum(q'(-logq'+logq))
  #this term is tricky because when q'=0 for a given species we get 0*log0, which r has difficulty calculating
  #to follow the convention that 0log0=0 we separate this case from cases where qd>0
  term3<-(1-P)*(sum(qdC[qdC>0]*(-log(qdC[qdC>0])- -log(qaC[qdC>0]))) #case 1 q'>0
                +sum(qdC[qdC==0]*(--log(qaC[qdC==0])))               #case 2 q'=0
  )
  
  
  
  
  pTerms<-c(term1,term2,term3)
  
  return(pTerms)
}


plotPriceEq<-function(qa,qd,alpha1,za,zd,zStar,P, YLAB="Trait value", YLIM="default") {
  
  #frequency in ancestral pop
  #frequency of decendants of ancestral pop
  #frequency of immigrants in species j
  #z trait value ancestors
  #trait value of decendants
  #trait value of immigrants
  #proportion of immigrants
  
  #first reformat to calculate trait change with "old species" by weeding out species only found among
  #decendants
  
  qaC<-qa[qa>0]
  qdC<-qd[qa>0]
  zaC<-za[qa>0]
  
  #first calculate terms involving non-immigrants
  wi=qdC/qaC                                  
  deltaZ<-zd-za
  
  #ignore "new species"
  deltaZ<-deltaZ[qa>0]
  
  par(mfrow=c(2,1), mar=c(1, 4, 1, 2) + 0.)
  
  ymax<-max(za,zd,zStar)
  ymin<-min(za,zd,zStar,-0.2)
  
  if(YLIM[1]=="default"){
    YLIM=c(ymin,ymax)
  }
  plot(0,1, type="n", xlim=c(0,1),ylim=YLIM,xlab="",ylab=YLAB)
  abline(h=0)

  #place line for proportion of immigrants
  abline(v=1-P, col="red")
  #draw a box representing proportion of species 1*its trait value
  
  
  
  #this calculates coordinates that are used several times in plots, note that the 1-P weights things by frequency of connected individuals
  b1x1<-0
  b1x2<-qa[1]*(1-P)
  deltab1x<-qd[1]*(1-P)
  deltaTx<-(qaC*wi)[1]*(1-P)
  
  b2x2<-(qa[1]+qa[2])*(1-P)
  deltaTxb<-(qa[1]+qa[2]-(qaC*wi)[2])*(1-P)
  
  quid<-qd[1]*(1-P)
  
  
 
  
  
  abline(v=b1x2,col="purple")
  abline(h=za[1],col="purple")
  
  #this represents the "ancestral" trait values
  polygon(c(b1x1,b1x1,b1x2,b1x2),y=c(za[1],0,0,za[1]),col="lightgrey")
  #draw a box representing effect of change in species 1's frequency
  polygon(c(deltab1x,deltab1x,b1x2,b1x2),
          y=c(za[1],0,0,za[1]),
          col=ifelse(qa[1]>qd[1],"#A6CEE3","#1F78B4"))
  
  #draw trait change
  polygon(c(b1x1,b1x1,deltaTx,deltaTx),
          y=c(za[1],za[1]+deltaZ[1],za[1]+deltaZ[1],za[1]),
          col=ifelse(deltaZ[1]<0,"#B2DF8A", "#33A02C"))  
  
  
  
  
  ########box diagram for species 2
  
  
  plot(0,1, type="n", ylim=YLIM,xlim=c(0,1),xlab="proportion of decendant community",ylab=YLAB,YLIM=c(0,1))
  abline(h=0)
  abline(v=b1x2,col="purple")
  abline(h=za[2],col="purple")
  
  #draw a box represeting proportion of species 2 times its trait value, but placed so that it abuts previous box
  polygon(c(b1x2,b1x2,b2x2,b2x2),y=c(za[2],0,0,za[2]),col="lightgrey")
  
  #change due to selection/drift species 2
  polygon(c(quid,quid,b1x2,b1x2),
          y=c(za[2],0,0,za[2]),
          col=ifelse(qa[1]<qd[1],"#A6CEE3","#1F78B4"))
  #trait change
  polygon(c(b2x2,b2x2,deltaTxb,deltaTxb),
          y=c(za[2],za[2]+deltaZ[2],za[2]+deltaZ[2],za[2]), 
          col=ifelse(deltaZ[2]<0,"#B2DF8A", "#33A02C"))
  
  mHeight<-abs(sum(alpha1*zStar)-sum(qa*za)) #difference in trait values between migrants and ancestors
  
  if(P>0){
  polygon(x=c(b2x2,b2x2,1,1),
          y=c(0,mHeight,mHeight,0),
          col="black")
  }
  
  return(NULL)
}


findFrankValues <- function(adMatrix) {
  #this function takes a matrix consisting of the species identity of each individual in an ancestral
  #and decendant community then returns the parameters required for franks version of the extended price equation
  #this function assumes that new individuals arrive only by speciation
  #this function assumes a fixed community size.
  
  #it is intended to pre-process for classicPriceExtended1
  
  
  comSize<-dim(adMatrix)[2]
  #table of abundance of each species at ancestral and decendant time step
  adTable<-data.frame(species=as.vector(adMatrix), generation=rep(c("a","d"),comSize))
  adTable<-table(adTable)
  Pa<-adTable[,1]/comSize
  
  #frequency of decendants of ancestral pop
  Pd<-adTable[,2]/comSize
  
  #the highest species number associated with ancestral community
  highSpANum<-max(as.numeric(names(Pa[Pa>0])),1)
  
  #Which decendants belong to old species
  dOldSp<-as.numeric(names(Pd))<=highSpANum
  PdOld<-Pd
  PdOld[!dOldSp]<-0
  PdOld<-PdOld/sum(PdOld)
  
  #frequency of new species
  PNew<-Pd
  PNew[dOldSp]<-0 #this is non normalized, useful for trait values for new species
  alpha<-PNew/sum(PNew) #this is normalized, as per frank's formula
  ProportionNew<-sum(PNew)
  
  FrankValues<-list(Pa=Pa,Pd=Pd,PdOld=PdOld,PNew=PNew,alpha=alpha,ProportionNew=ProportionNew)
  
  
  return(FrankValues)
}


#here are two functions to compute population growth. 


annualPlantGrowth<-function(N1,N2,r1,r2, stochastic=FALSE){
  #density of species 1
  N1P<-r1*N1/(1+a11*N1+a12*N2)
  
  #density of species 2
  N2P<-r2*N2/(1+a21*N1+a22*N2)
  
  if(stochastic==TRUE){
    N1P<-rpois(1,N1P)
    N2P<-rpois(1,N2P)
  }
  return(c(N1P,N2P))
}


rickerCompetition<-function(N1,N2, r1,r2,stochastic=FALSE){
  #density of species 1
  N1P<-N1*exp(r1-a11*N1-a12*N2)
  
  #density of species 2
  N2P<-N2*exp(r2-a22*N2-a21*N1)
  
  if(stochastic==TRUE){
    N1P<-rpois(1,N1P)
    N2P<-rpois(1,N2P)
  }
  return(c(N1P,N2P))
}


#exeperiment with alternative way to add stochasticity
rickerCompetition1<-function(N1,N2,stochastic=FALSE){
  
  
  if(stochastic==TRUE){
    #density of species 1
    N1P<-N1*exp(r1-a11*N1-a12*N2+rnorm(1,0,.02))
    
    #density of species 2
    N2P<-N2*exp(r2-a22*N2-a21*N1+rnorm(1,0,.02))
  }
  else{
    #density of species 1
    N1P<-N1*exp(r1-a11*N1-a12*N2)
    
    #density of species 2
    N2P<-N2*exp(r2-a22*N2-a21*N1)
  }
  
  return(c(N1P,N2P))
}


#exeperiment with alternative way to add stochasticity
rickerCompetition2<-function(N1,N2,r1,r2,stochastic=FALSE){
  
  
  if(stochastic==TRUE){
    #density of species 1
    N1P<-N1*exp(r1-a11*N1-a12*N2+rnorm(1,0,.02))
    
    #density of species 2
    N2P<-N2*exp(r2-a22*N2-a21*N1+rnorm(1,0,.02))
  }
  else{
    #density of species 1
    N1P<-N1*exp(r1-a11*N1-a12*N2)
    
    #density of species 2
    N2P<-N2*exp(r2-a22*N2-a21*N1)
  }
  
  return(c(N1P,N2P))
}


#exeperiment with alternative way to add stochasticity
rickerCompetition3species<-function(N1,N2,N3,stochastic=FALSE){
  
  
  if(stochastic==TRUE){
    #density of species 1
    N1P<-N1*exp(r1-a11*N1-a12*N2-a13*N3+rnorm(1,0,.02))
    
    #density of species 2
    N2P<-N2*exp(r2-a22*N2-a21*N1-a23*N3+rnorm(1,0,.02))
    
    N3P<-N3*exp(r3-a33*N3-a31*N1-a32*N2+rnorm(1,0,.02))
    
  }
  else{
    N1P<-N1*exp(r1-a11*N1-a12*N2-a13*N3)
    
    #density of species 2
    N2P<-N2*exp(r2-a22*N2-a21*N1-a23*N3)
    
    N3P<-N3*exp(r3-a33*N3-a31*N1-a32*N2)
    
  }
  
  return(c(N1P,N2P,N3P))
}


calculateDZ<-function(qa,qd,za,zd){
  
  #This function is designed to compute change within any species. 
  #qa is the frequency of all genotypes in the ancstral genration
  #qd is same for decendant generation
  #za and zd are traits assocaited with ancestral and decendant generations respectively.
  #as of now, this funciton is not fancy enough to deal with immigration, but it probably could be. 
  
  z1<-sum((qd-qa)*za) #change in species level trait value due to genotype level selection
  z2<-sum(qd*(zd-za)) #change in species level trait value due to "genotype level" trait change
  
  #by term1 I mean the the total change at the species level due to seleciton at the genotype level, this is a factor
  #times deltaZ, looking at my notes i believe it should be qd times this factor, though re-check with whiteboard. 
  
  #term1<-NA
  
  #by term1 2mean the the total change at the species level due to trait change at the genotype level, this is a factor
  #times deltaZ, looking at my notes i believe it should be qd times this factor, though re-check with whiteboard. 
  #term2<-NA
  
  return(c(z1,z2))
}


LogNegCon<-function(x){
  #normally log(0) is undefined. R by default returns -Inf. HOwever, In the calculations we do downstream we do vectorized
  #calculatiosn where,
  #log(0) occurs. but one of two things happens, either log(0) is calculated because of the way vecotrs are produced here
  #but not counted in a sum, or log(0) appears in an operation like 0log0. By convention (Cover and Thomas elements of information theory).
  #0log0=0. A painless way to deal with both cases  in the code i have is to set log0=0. This will not work for other applications.
  x<- -log(x)
  x[x==Inf]<-0
  return(x)
}
```
